# Patient iPSC-Derived Cartilage Organoids Reveal Defective ECM Deposition and Altered Chondrogenic Trajectory in Saul-Wilson Syndrome

**DOI:** 10.64898/2026.04.10.717608

**Authors:** Sonal Mahajan, Sara Ancel, Giuliana Ascone, Rajdeep Kaur, Juancarlos Torres, Rabi Murad, Yu Xin Wang, Carlos R. Ferreira, Hudson H. Freeze

**Author notes:** Co-senior author (Sonal Mahajan, Hudson H. Freeze).

## Abstract

Saul-Wilson syndrome (SWS) is a skeletal dysplasia characterized by primordial dwarfism and progeroid features caused by a recurrent dominant COG4 variant (p.G516R). We previously showed that this mutation accelerates Golgi retrograde trafficking and disrupts glycosylation of the proteoglycan decorin, while zebrafish models revealed defects in chondrocyte elongation and intercalation. We have also shown that the SW1353 chondrosarcoma cells carrying the SWS variant exhibit reduced secretion of extracellular matrix (ECM) components. While these results indicate a critical function of COG4 in Golgi processing, the developmental process leading to skeletal dysplasia in SWS patients remains unknown. Here, we generated patient-derived iPSC cartilage organoids (SWS organoids), modeling early human chondrogenesis. SWS organoids failed to produce cartilage structures and displayed poor expression of chondrogenic markers. Time-course RNA-seq analysis of the chondrogenic process revealed reduced activation of gene networks involved in skeletal development, ECM organization, ossification, and glycosaminoglycan metabolism. Spatial multiomic analysis of protein and glycosylation by CODEX and GLYPH imaging revealed an altered chondrogenic trajectory, persistence of mesenchymal states, global glycosylation changes, and reduced deposition of chondroitin sulfate proteoglycans. These results indicate that the COG4 mutation disrupts ECM glycosylation and chondrogenic commitment, and that SWS organoids model early defects in cartilage formation underlies impaired skeletal growth in SWS.

**Highlights:**

1. Patient iPSC-derived cartilage organoids model development defects in Saul–Wilson syndrome
2. SWS organoids show defective extracellular matrix deposition and attenuated chondrogenic gene expression
3. Glycan profiling reveals global glycosylation defects and deficient proteoglycan GAG chains
4. An early developmental impairment in chondrogenesis alters skeletal formation in Saul–Wilson syndrome

## Introduction

Saul–Wilson syndrome (SWS) is a rare genetic disorder caused by a recurrent heterozygous *de novo* variants in COG4 (MIM: 606976, NM_015386.2:c.1546G>A or c.1546G>C; p.Gly516Arg), a Golgi-localized protein that is part of the conserved oligomeric Golgi (COG) trafficking complex (C. R. Ferreira et al., 2018; Xia et al., 2022). The COG complex consists of eight subunits (COG1–8) and plays a central role in vesicle tethering and Golgi homeostasis, thereby regulating both retrograde and anterograde vesicular trafficking (Ungar et al., 2002).

While biallelic variants in COG4 and other COG subunits cause Congenital Disorders of Glycosylation (CDG), characterized by abnormal N-glycosylation and intellectual and neurological impairments (D’Souza et al., 2020; Ng et al., 2011; Reynders et al., 2009), the recurrent p.G516R variant results in a clinically distinct disorder primarily affecting skeletal development (C. R. Ferreira et al., 2018).

SWS is characterized by primordial dwarfism, severe prenatal and postnatal growth restriction, and distinctive craniofacial and radiographic features (C. Ferreira, 2020; C. R. Ferreira et al., 2018, 2020). Additional clinical manifestations include cataracts, retinal dystrophy, hearing loss, micrognathia, brachydactyly, and skeletal abnormalities affecting both axial and appendicular elements (C. Ferreira, 2020; C. R. Ferreira et al., 2018, 2020). To date, only sixteen individuals with SWS have been described worldwide, underscoring the extreme rarity of this condition.

Previous studies using patient fibroblasts and engineered SW1353 chondrosarcoma cell lines harboring the COG4 p.G516R variant identified accelerated Golgi retrograde trafficking and abnormal glycosylation of decorin, a key extracellular matrix (ECM) proteoglycan involved in cartilage organization (C. R. Ferreira et al., 2018; Xia et al., 2022). Additionally, both SW1353 and HEK293T cells with the mutation showed subtle, cell-type dependent changes in N-glycans (Xia et al., 2022). Complementary studies in zebrafish expressing the SWS variant demonstrated reduced body size, abnormal fin morphology, and disrupted chondrocyte elongation and intercalation, accompanied by altered WNT4 signaling during embryogenesis (Xia et al., 2021). Despite these advances, existing experimental systems have provided only partial insight into disease mechanisms. In particular, the SW1353 chondrosarcoma model showed variability between clones and limited capacity for sustained chondrogenesis due to its transformed and terminally differentiated nature. Furthermore, mouse models fail to recapitulate the heterozygous skeletal phenotype observed in patients, highlighting the need for lineage-relevant human systems to investigate SWS pathogenesis.

Here, we address these limitations by establishing patient-derived induced pluripotent stem cell (iPSC) to cartilage organoids as a platform to model SWS. This system enables the study of COG4-associated defects in a human genetic context while preserving developmental relevance and chondrogenic capacity. By leveraging this system, we interrogate cell-type-specific glycosylation defects, extracellular matrix organization, and disease-relevant signaling pathways in a controlled and reproducible manner.

Importantly, using this model, we reveal defects in extracellular matrix organization associated with impaired glycosylation and proteoglycan maturation, offering novel insight into how Golgi dysfunction leads to skeletal abnormalities in SWS. Together, our findings establish a disease-relevant human model that overcomes key limitations of prior systems and provide a foundation for mechanistic studies and future therapeutic exploration.

## Results

### Mouse models fail to reproduce the human skeletal phenotype

To assess the in vivo consequences of the *COG4^+/G516R^* variant, we generated knock-in mice carrying the orthologous variant (*Cog4^+/G512R^*). Heterozygous *Cog4^+/G512R^* mice displayed normal skeletal architecture and bone mineral density, whereas homozygous *Cog4^G512R/G512R^* mutants were embryonic lethal and exhibited severe defects in bone growth and mineralization (Figures S1–S2). These findings indicate that heterozygous expression of the variant does not disrupt murine skeletal development, whereas homozygous expression leads to severe skeletal defects, with in utero lethality, suggesting species-specific sensitivity to COG4 dosage.

### Patient iPSC-derived mesenchymal progenitors retain the Golgi trafficking defect

Given the absence of a skeletal phenotype in heterozygous mice, we established patient-derived induced pluripotent stem cell (iPSC) based cartilage models to investigate the impact of the *COG4^+/G516R^* variant in a human developmental context. Control and patient fibroblast-derived iPSCs were differentiated into mesenchymal progenitor cells (MPCs) and subsequently into chondrocytes (Figure 1A). After 13 days of induction, both control and patient lines expressed canonical mesenchymal markers CD90, CD73, and CD146, while lacking the hematopoietic marker CD45 (Figure S3A).

**Figure 1.**
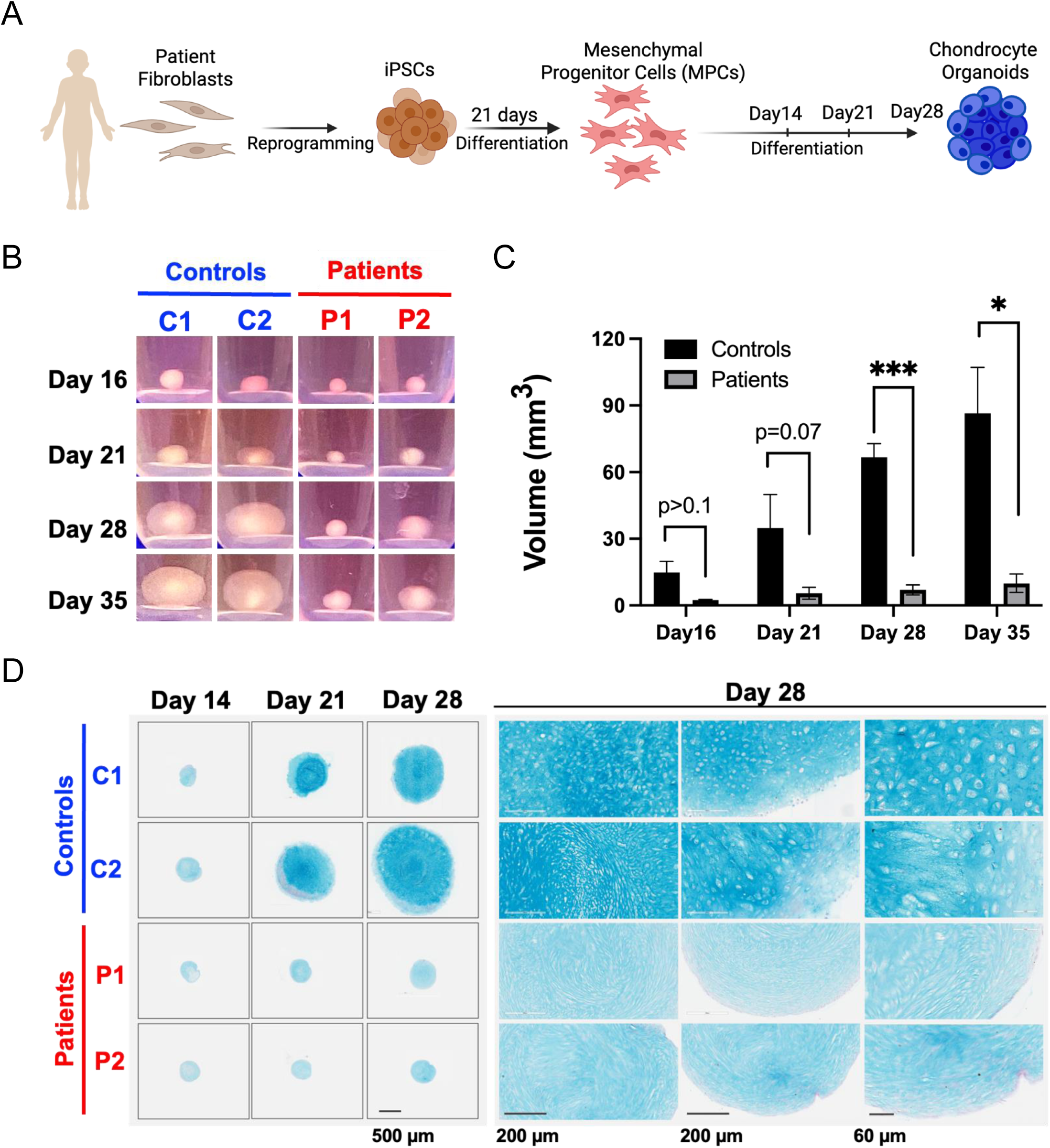
Patient iPSC-derived cartilage organoids exhibit reduced size and less deposition of acidic polysaccharides. A. Experimental overview for generating patient iPSC-derived cartilage organoids. B. Representative images of cartilage organoids at different time points during differentiation. Images are representative of three independent experiments with similar results. C. Quantification of cartilage organoid volume in control and patient lines over time. P values represent statistical significance determined by two-way ANOVA (n = 3). D. Alcian blue staining of cartilage organoids showing deposition of acidic polysaccharides. Experiments were performed in triplicate with comparable results.

To confirm that MPCs retained the SWS cellular phenotype, we examined Golgi trafficking dynamics using a brefeldin A (BFA) - induced retrograde transport assay. Imaging flow cytometry revealed accelerated Golgi fragmentation in patient MPCs compared with controls following BFA treatment (Figs. S3B–S3D), consistent with previously reported defects in COG4 mutant cells. These findings indicate that patient-derived MPCs retain the characteristic Golgi trafficking phenotype of SWS and provide a suitable platform for downstream chondrogenic differentiation.

### SWS cartilage organoids exhibit impaired growth and extracellular matrix deposition

Chondrogenic differentiation of MPCs was performed using a high-density 3D culture system (Guzzo et al., 2013; Koyama et al., 2012; Ullah et al., 2012). Control and SWS patient iPSC- derived organoids (SWS organoids) displayed comparable growth until approximately Day 15. However, control organoids subsequently underwent substantial volumetric expansion, whereas SWS organoids failed to increase in size (Figure 1B). By Days 21–35, control organoids were 8–12-fold larger than SWS organoids (Figure 1B, 1C), consistent with extensive extracellular matrix deposition during cartilage maturation. In contrast, SWS organoids remained compact and lacked the matrix-rich architecture observed in controls.

Alcian blue staining confirmed the deposition of sulfated proteoglycan at late time points in control organoids, consistent with robust chondrogenesis (Figure 1D). Consistent with their reduced size, SWS organoids exhibited markedly weaker Alcian blue staining compared with controls, indicating diminished cartilage matrix production (Figure 1D).

Morphological differences were also evident. At Day 28, control organoids displayed a growth plate–like organization, with flattened cells at the periphery, rounded chondrocyte-like cells in the intermediate zone, and densely packed cells toward the core (Figure 1D). In contrast, SWS organoids lacked this organized architecture and were composed predominantly of elongated, spindle-shaped mesenchymal-like cells (Figure 1D).

### SWS organoids fail to activate chondrogenic transcriptional programs

To assess chondrogenic differentiation, we measured mRNA levels of early (*SOX9, ACAN, COL2A1*) and late (*COL10A1, IHH*) chondrocyte markers. Control organoids exhibited progressive induction of early markers between MPCs and Day 28, followed by a decline at Day 35 consistent with hypertrophic maturation (Figure 2A). In contrast, SWS organoids showed minimal induction of these markers, with expression levels remaining 3–4-fold lower than controls. Late chondrocyte markers, including *IHH* and *COL10A1*, were robustly expressed in control organoids but were nearly absent in SWS organoids (Figure 2A). These findings indicate impaired activation of chondrogenic transcriptional programs in SWS organoids.

**Figure 2.**
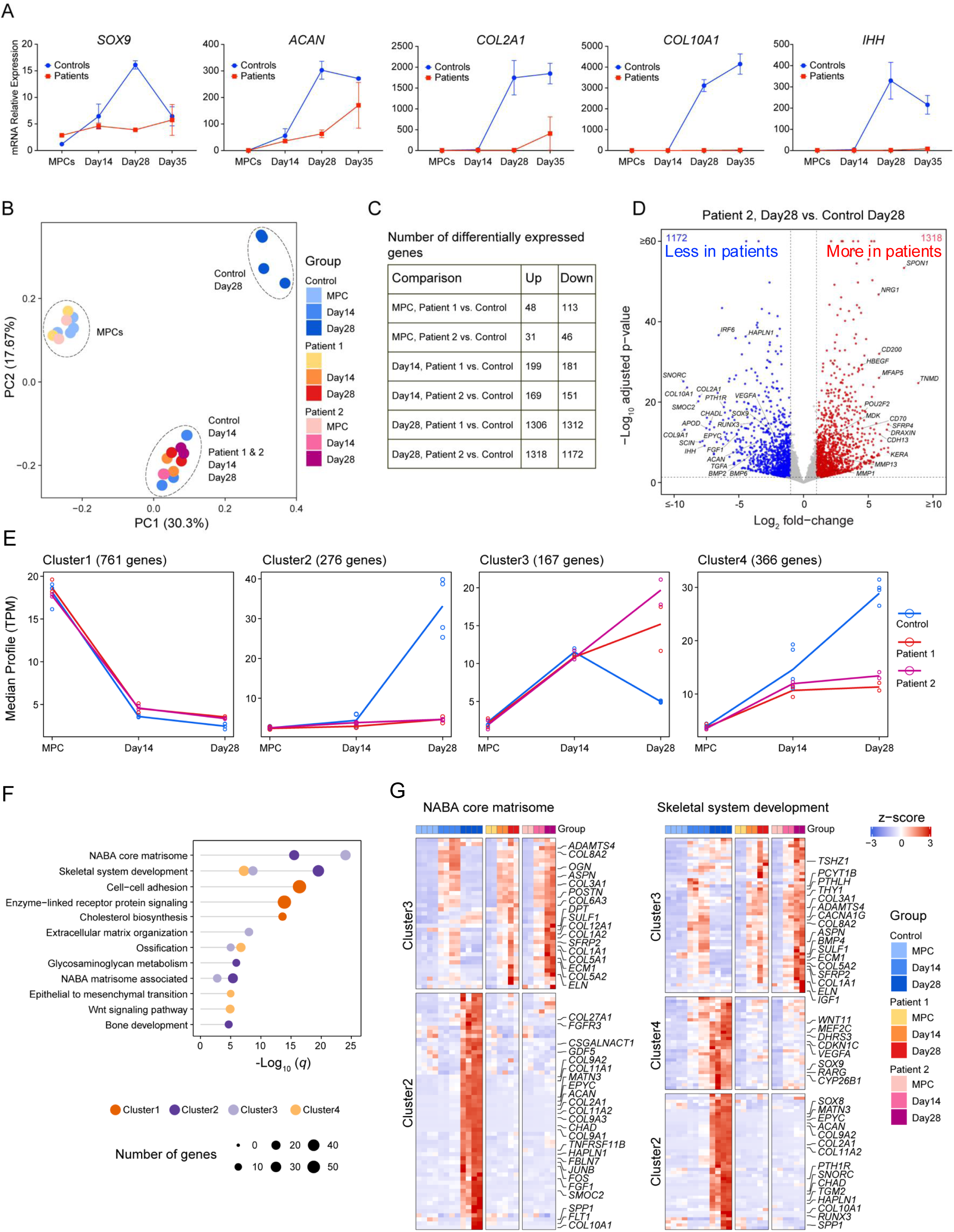
SWS organoids demonstrate deficient expression of chondrogenic markers at the transcriptional level. A. qPCR analysis of chondrocyte differentiation markers. Error bars represent the standard deviation of two biological replicates. B. Principal component analysis (PCA) of mRNA-seq data showing variation between samples. Control and patient samples were analyzed in duplicates. For the analysis, all four control time points were combined and compared with duplicate samples from each patient. C. Table showing the number of genes that are differentially expressed between control and patient samples. D. Volcano plot showing differentially expressed genes in control and SWS organoids. Blue dots represent genes with higher expression in controls, while red dots represent genes with higher expression in patient samples. The y-axis denotes −log10 adjusted p-values, and the x-axis shows log2 fold-change values. E. Gene expression profile clustering using the maSigPro R package to identify groups of genes with significant expression patterns during chondrogenic differentiation. A total of nine clusters with distinct temporal profiles were identified; the figure shows the four most significant and physiologically relevant clusters. (All clusters are shown in in the Supplementary Figure S4). F. Bar chart showing Metascape enrichment analysis of ontology categories (GO and KEGG terms) for the selected clusters. G. Heatmaps showing differential expression patterns for enriched pathways, including skeletal system development and the NABA Core Matrisome, across different time points.

To define transcriptional programs underlying the differentiation defect, we performed RNA sequencing (RNA-seq) across multiple stages of organoid formation. Principal component analysis revealed clear separation of control MPCs, Day 14, and Day 28 organoids, reflecting progressive differentiation (Figure 2B). In contrast, SWS Day 28 organoids clustered with Day 14 organoids, indicating stalled developmental progression.

Differential gene expression analysis further highlighted the divergence between control and SWS organoids during later stages of differentiation. The largest transcriptional differences occurred at Day 28, with 1306 and 1318 genes significantly upregulated in Patient 1 and Patient 2 compared with controls, respectively, and 1312 and 1172 genes significantly downregulated (Figure 2C). In contrast, fewer than 200 genes were differentially expressed between control and SWS at the MPC stage and Day 14 organoids (Figure 2C), consistent with the similar transcriptional profiles observed at early stages.

SWS organoids showed marked downregulation of key chondrogenic and cartilage matrix genes, including *SOX9, COL2A1, COL9A1, ACAN, EPYC, HAPLN1*, and *IHH*, together with reduced expression of regulators of chondrocyte maturation and skeletal development such as *PTH1R, BMP2/6,* and *RUNX3* (Figure 2D). In contrast, genes associated with matrix remodeling or fibroblastic and non-chondrogenic lineages were upregulated, including *MMP1, MMP13, SPON1, SFRP4, TNMD,* and *MFAP5* (Figure 2D), indicating impaired activation of cartilage-specific transcriptional programs and a shift toward alternative mesenchymal trajectories.

Time-course gene expression analysis using the maSigPro algorithm identified nine gene clusters with distinct expression dynamics during differentiation (Figure S4), with representative clusters shown in Figure 2E. Cluster 1 (761 genes) comprised of genes progressively downregulated during differentiation in both control and SWS organoids and was enriched for pathways related to cell–cell adhesion, receptor signaling, and cholesterol biosynthesis (Figures 2E–2F).

In contrast, Clusters 2 (276 genes) and 4 (366 genes) contained genes that were strongly induced in control organoids at later stages (Day 14–Day 28) but failed to be activated in SWS organoids (Figure 2E). Together with Cluster 3 (167 genes), these clusters were enriched for pathways related to the NABA core matrisome, skeletal system development, extracellular matrix organization, ossification, and glycosaminoglycan metabolism (Figure 2F). Control organoids showed robust induction of chondrogenic and ossification-associated genes, including *SOX9, ACAN, COL2A1, SNORC, SMOC2, CHADL, CHAD, IHH, SPP1*, multiple *BMPs*, and collagen genes (Figure 2G).

Conversely, SWS organoids failed to activate these programs and instead maintained expression of genes associated with mesenchymal or fibroblastic states, including *THY1*, *COL1A1, COL1A2, COL3A1, SPARC, POSTN, BMP4,* and *FGFR2*, consistent with persistence of a mesenchymal stromal/progenitor-like identity rather than progression to terminal chondrocytes.

### Spatial multi-omic analysis of cartilage organoids reveals stalled chondrogenic differentiation and persistence of mesenchymal states in SWS

To determine whether transcriptional differences observed during chondrogenesis were reflected at the protein level, we performed spatial multiomic analysis using highly multiplexed immunofluorescence imaging of cellular and ECM proteins (CODEX) (Singla et al., 2025) as well as glycosylation via Glycosylation Landscape anaLysis by Probe Hybridization (GLYPH) with a custom panel DNA-barcoded glycan-binding lectins we developed for this study. This enabled us to spatially resolve cell populations, their surrounding ECM, and glycosylation profiles within organoid sections. In total, 46 markers were simultaneously visualized (Supplementary Table S1), including markers of chondrocytes, extracellular matrix (ECM), mesenchymal progenitors, and glycan-binding lectins. Antibodies and lectins that did not perform reliably were excluded from further analysis.

In control organoids, SOX9⁺ cells first appeared at Day 14 and increased progressively at Days 21 and 28. In contrast, SOX9 expression was largely absent in SWS organoids (Figure 3A, 3B). Consistent with this pattern, the cartilage matrix proteins aggrecan and COL2A1 accumulated over time in control organoids but remained markedly reduced in SWS organoids (Figure 3A, 3B). Markers associated with late chondrocyte maturation and hypertrophy including RUNX2, COLX, IHH, MMP13, and CD200 emerged at Day 21 or Day 28 in control organoids but showed minimal or no expression in SWS organoids. Representative images for RUNX2 expression and the quantification is shown in Figure 3A and 3B.

**Figure 3:**
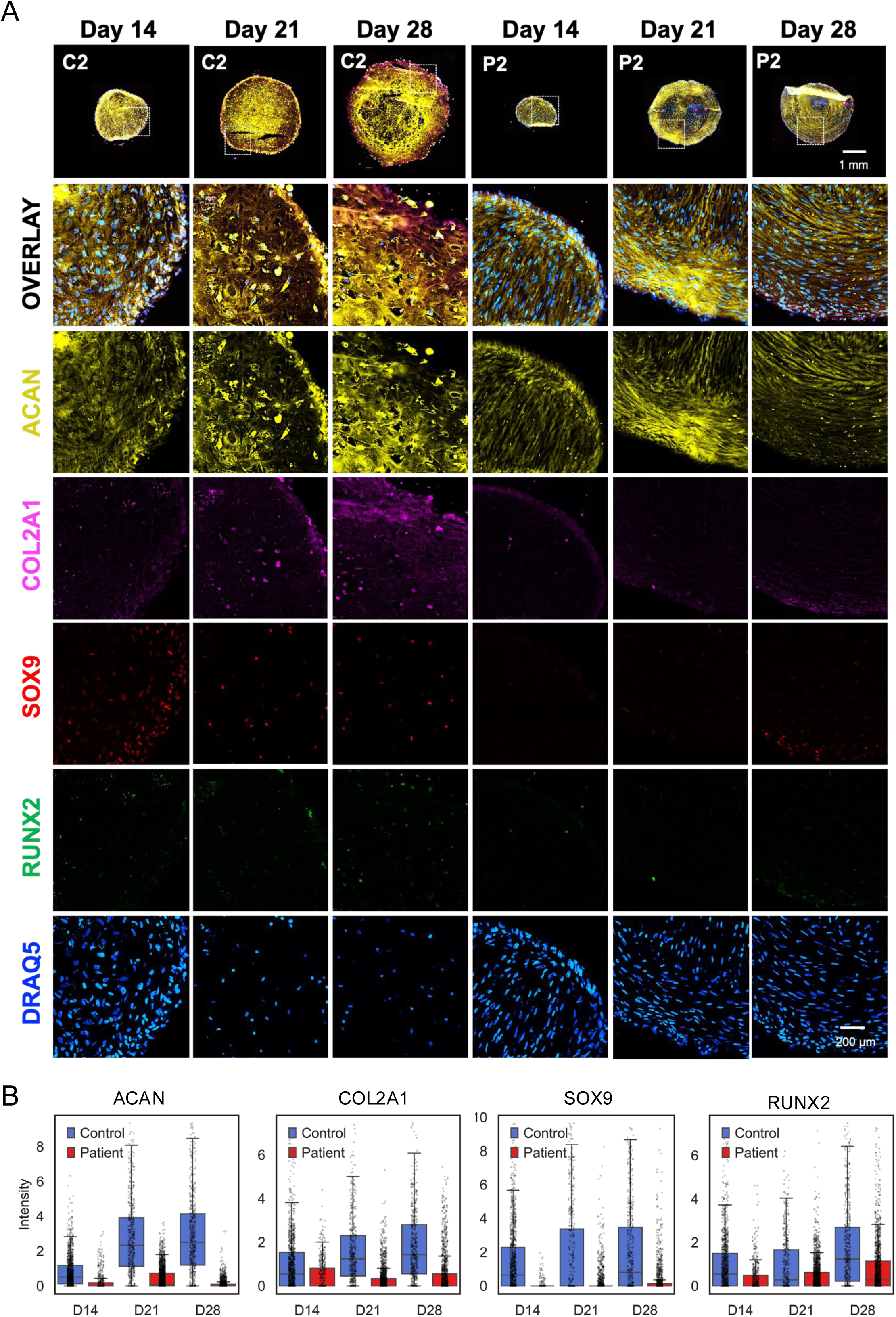
CODEX staining of canonical chondrogenesis markers reveals impaired chondrocyte differentiation in SWS organoids. A. Representative images of organoid sections highlighting the following markers: DRAQ5, SOX9, Aggrecan, COL2A1, and RUNX2. Images are representative of three independent experiments with consistent results. B. Quantification of fluorescence intensity per cell, illustrating the distribution of marker expression within individual organoids, derived from CODEX imaging data.

Conversely, mesenchymal-associated markers (COL1A1, CD44, and CD9) progressively declined in control organoids as progenitors differentiated into chondrocytes (Figure 4A, 4B). In SWS organoids, however, these markers remained persistently expressed at later time points, indicating a failure to exit the mesenchymal state and suggesting a block in chondrogenic differentiation.

**Figure 4:**
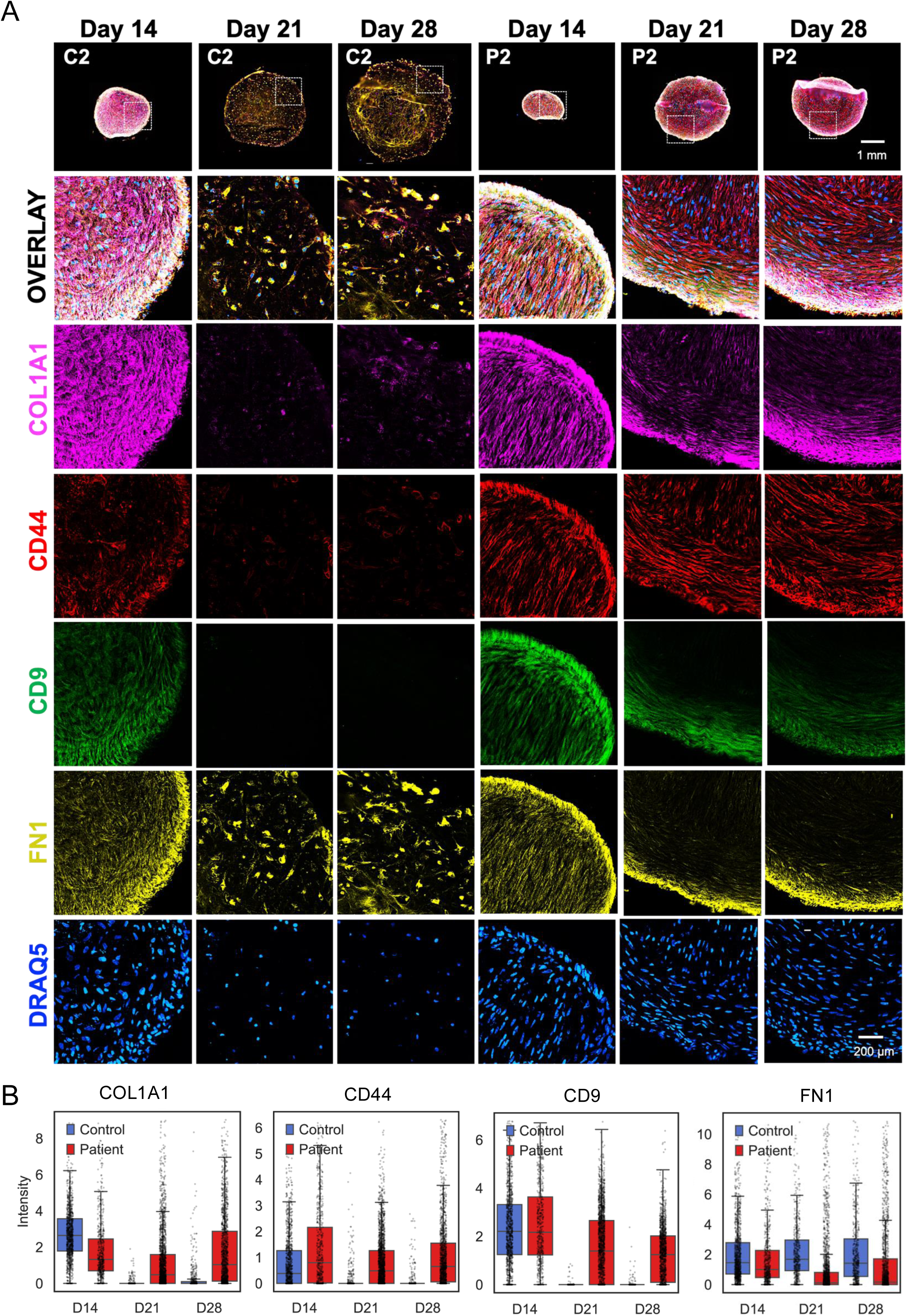
CODEX staining reveals enrichment of mesenchymal-like cell populations in SWS organoids. A. Representative images of organoid sections imaged using CODEX, highlighting the following markers: COL1A1, CD44, CD9, and Fibronectin. Images are representative of three independent experiments with consistent results. B. Quantification of fluorescence intensity per cell of the indicated markers, illustrating the distribution of marker expression within individual organoids, derived from CODEX imaging data.

To further define cell populations, unsupervised clustering of single-cell protein expression profiles from CODEX generated a uniform manifold approximation and projection (UMAP) representation comprising ten distinct clusters across all conditions (Figure 5A). Clusters were annotated as distinct cellular states based on marker co-expression patterns (Figure 5B).

**Figure 5:**
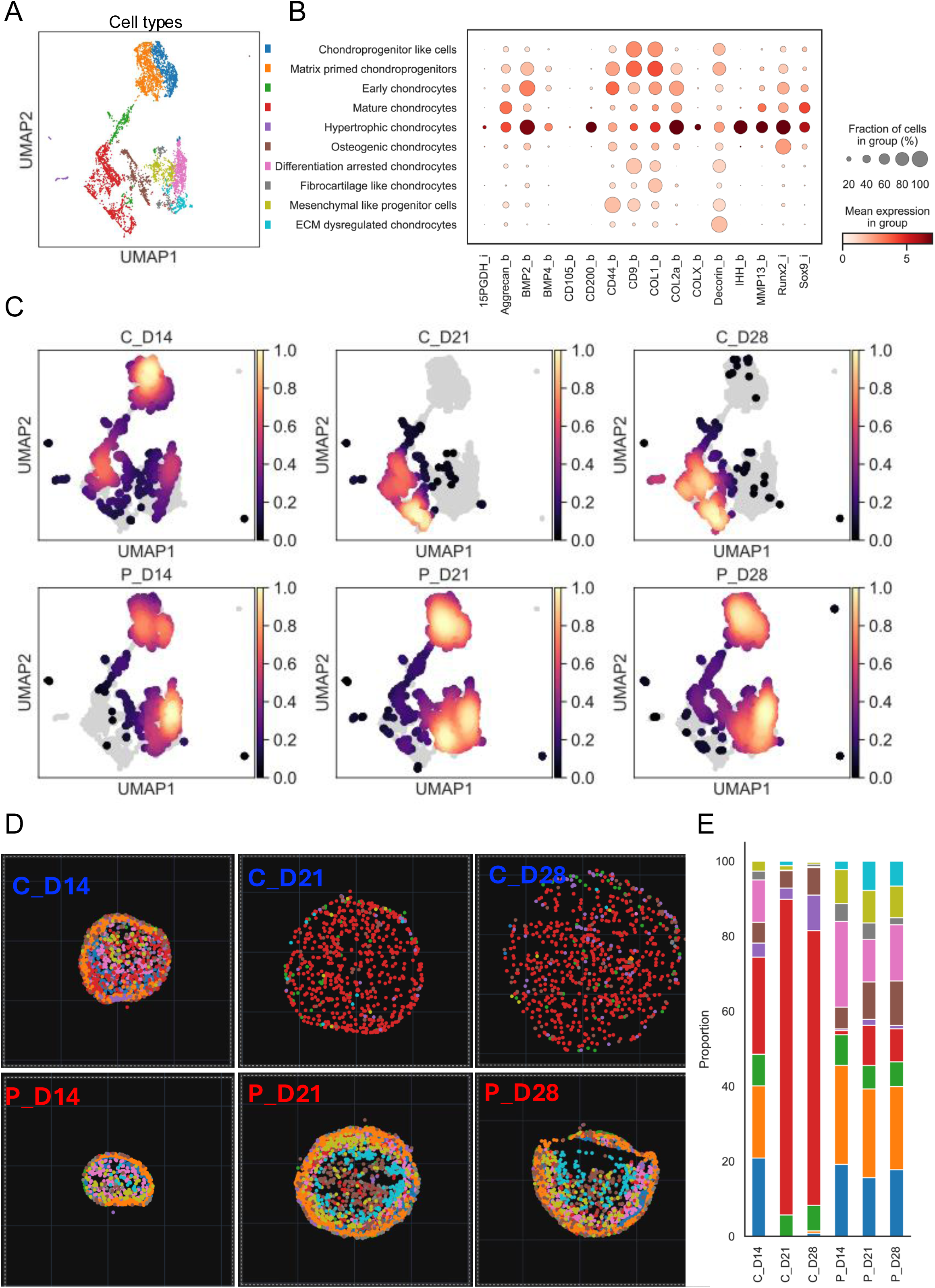
Spatial profiling by multiplexed CODEX staining reveals enrichment of mesenchymal-like cell populations and a developmental block during chondrogenesis in SWS organoids. A. UMAP embedding of 17,866 cells from control and SWS organoids, colored by annotated cell types. The color scheme is consistent across panels B, D, and E. B. Dot plot showing scaled expression of marker proteins across identified cell clusters, defining distinct cellular identities. Dot size represents the percentage of cells within each cluster expressing a given marker, while color intensity indicates the average expression level. C. Kernel density estimates of cell distribution in UMAP space for each condition (Control vs SWS). Densities are normalized within each condition to highlight relative enrichment patterns along chondrogenic trajectories. D. Spatial maps showing the distribution of cell types based on cluster annotations projected in spatial coordinates for each condition, corresponding to distinct stages of chondrogenesis. E. Graph showing the proportions of different cell types in control and SWS organoids across the analyzed time points.

Kernel density estimates of cell distribution across conditions revealed distinct enrichment patterns in UMAP space, with control organoids preferentially occupying regions corresponding to mature chondrocyte states, whereas SWS organoids remained enriched in mesenchymal-like and intermediate populations (Figure 5C). Spatial maps were generated to visualize the distribution of various cell populations within organoid sections (Figure 5D).

At Day 14, control organoids exhibited a heterogeneous cellular composition, comprising CD9⁺ and COL1⁺ chondroprogenitor-like cells (∼20%), ACAN⁺ and COL2A⁺ matrix-primed chondroprogenitors (∼20%), SOX9⁺ and COL2A⁺ early chondrocytes (∼8–10%), and mature chondrocytes (∼25%). The remaining cells included IHH and MMP13⁺ hypertrophic cells as well as CD44⁺ mesenchymal-like populations (Figure 5D, 5E). Spatially, early progenitors were predominantly localized to the organoid periphery, whereas more differentiated chondrocytes were enriched toward the core, indicating a spatially organized maturation process. By Day 21, control organoids were largely composed of mature chondrocytes (∼85%), with a small fraction of hypertrophic chondrocytes (3–4%). Hypertrophic cells increased modestly by Day 28 (∼10%), consistent with progressive terminal maturation (Figure 5D, 5E).

In contrast, SWS organoids displayed an immature cellular composition as early as Day 14, characterized by an increased presence of mesenchymal-like progenitors (∼10%) and chondroprogenitors (∼20%) that retained progenitor markers such as CD9 but did not robustly upregulate SOX9, COL2A1, or aggrecan. This early-stage profile persisted through Days 21 and 28. At later time points, SWS organoids were dominated by chondroprogenitor-like (∼25%), COL1⁺ fibrocartilage-like (∼20%), and CD44⁺ mesenchymal-like (∼20%) populations, with only limited emergence of mature chondrocytes (5–10%) and RUNX2⁺ osteogenic or IHH and MMP13⁺ hypertrophic chondrocytes (5–10%). A subset of cells (5–8%) was classified as ECM-dysregulated chondrocytes based on decorin overexpression. Collectively, these findings indicate a stalled differentiation trajectory and impaired progression toward terminal chondrocyte maturation (Figure 5D, 5E).

Quantification of nuclei per organoid revealed no further increase in cell number in control organoids at Days 21 and 28, consistent with cell cycle exit accompanying differentiation (Figure S5A, S5B). A reduction in average nuclear intensity and decreased nuclear-to-cytoplasmic ratio further supported terminal differentiation (Figure S5C, S5D). In contrast, SWS organoids showed increasing nuclear counts from Day14 to Day21 and Day28, and higher nuclear-to-cytoplasmic ratios at later stages, consistent with sustained proliferation or incomplete differentiation (Figure S5A, S5B, S5C).

Spatial analysis of nuclear distribution relative to the organoid centroid revealed dispersed nuclei within an expanded ECM in control organoids, whereas SWS organoids exhibited densely packed nuclei and limited ECM expansion (Figure S5E). Notably, the increase in organoid volume in controls was driven primarily by ECM deposition rather than increased cell number, whereas SWS organoids showed restricted volumetric growth, consistent with impaired matrix production.

### GLYPH reveals global glycosylation defects and impaired proteoglycan modification in SWS organoids

Immunostaining of organoid sections with lectins from the GLYPH panel revealed distinct glycosylation patterns between control and patient samples (Figure 6A). Rather than reflecting differences in overall staining intensity alone, individual lectins labeled distinct cellular and extracellular structures in control versus SWS organoids, indicating widespread alterations in glycosylation of cell-associated and secreted proteins. Representative lectin staining patterns are shown in Figure 6A.

**Figure 6:**
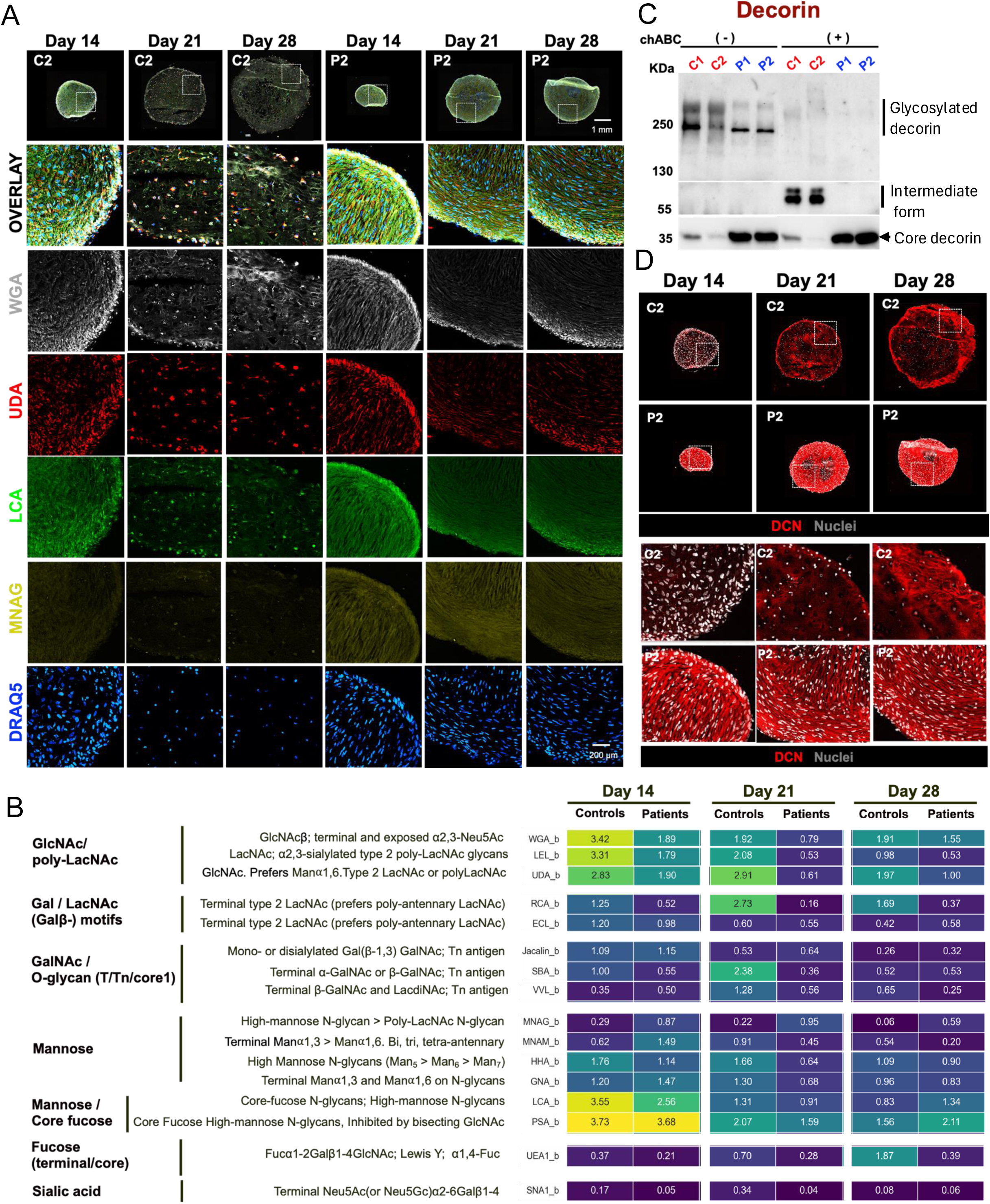
GLYPH imaging reveals altered glycosylation and deficient Glycoaminoglycans (GAG) in chondroitin sulfate proteoglycans in SWS organoids. A. Representative images of organoid sections imaged using CODEX, highlighting the following lectins: WGA, MNAG, UDA, and LCA, along with the nuclear marker DRAQ5. Images are representative of three independent experiments with similar results. B. Heatmap showing the coefficient of mean signal intensity per nucleus for each lectin in the CODEX panel across the indicated conditions. C. Western Blot analysis of Decorin showing core protein and the glycosylated forms of Decorin in the conditioned medium from Day 28 control and SWS organoids, before and after Chondroitinase ABC treatment. D. Representative images of organoid sections highlighting the distinct patterns of Decorin staining in control vs SWS organoids.

Lectin staining was quantified by measuring mean fluorescence intensity (MFI) per nucleus using AI-assisted analysis of GLYPH-stained images (Figure 6B). Quantitative comparisons were restricted to Day 14, when control and SWS organoids exhibit comparable cellular compositions. At later stages (Days 21 and 28), divergent differentiation states between control and SWS organoids precluded direct comparison due to cell-type–dependent glycosylation differences.

At Day 14, lectins recognizing GlcNAc/poly-LacNAc motifs, Gal/LacNAc motifs, GalNAc/O-glycan motifs, and sialic acid exhibited 2–3-fold higher signal in control organoids (Figure 6B). In contrast, lectins MNAG (Gal-binding) and MNAM (predominantly recognizing high-mannose structures) showed 2.5–3-fold higher signal in SWS organoids (Figure 6B). Lectins recognizing core fucosylation (LCA and PSA) showed no substantial differences between control and patient samples (Figure 6B).

Previous cellular models of SWS have demonstrated altered glycosylation of the proteoglycan decorin (C. R. Ferreira et al., 2018; Xia et al., 2022). To determine whether similar defects were present in organoids, conditioned medium from Day 28 organoids was analyzed by Western blot to distinguish glycosylated versus core decorin species. Patient samples exhibited a higher proportion of core decorin and reduced levels of higher–molecular weight glycosylated forms, which collapsed to the core protein following chondroitinase ABC (chABC) digestion (Figure 6C). In contrast, approximately 80% of decorin in control organoids migrated as higher–molecular weight glycosylated species and shifted to an intermediate form after chABC treatment (Figure 6C), indicating defective decorin glycosylation in SWS organoids.

We next examined aggrecan, another major chondroitin sulfate proteoglycan. In SWS organoids, aggrecan signal was faint and migrated slightly higher than in controls, suggesting altered glycosylation (Figure S6A). Following chABC treatment, an additional higher–molecular weight band appeared in control organoids, likely reflecting charge-related migration changes, but this band was absent in SWS organoids (Figure S6A).

Immunofluorescence analysis further revealed differences in decorin localization between control and SWS organoids (Figure 6D). In control samples, decorin was predominantly intracellular at Day 14, followed by increased expression and progressive accumulation within the extracellular matrix at Days 21 and 28. In contrast, SWS organoids displayed a distinct pattern in which decorin appeared more extracellular from early time points, potentially reflecting premature secretion of an underglycosylated form into the matrix (Figure 6D).

To directly assess chondroitin sulfate glycosaminoglycan (GAG) composition, we performed disaccharide analysis. SWS organoids exhibited an approximately 90% reduction in total chondroitin sulfate disaccharides, accompanied by 7- to 12-fold decreases in 4-O-SO₃ and 6-O-SO₃ chondroitin sulfate disaccharides, respectively (Figure S6B–E). Together, these findings demonstrate a profound disruption of proteoglycan glycosylation and GAG modification in SWS organoids.

## Discussion

In this study, we employed iPSC-derived cartilage organoids as a patient-specific and physiologically relevant model to investigate the pathology of Saul–Wilson syndrome. In our previous work using SW1353 chondrosarcoma cells, we observed impaired formation of chondrocyte spheroids and reduced secretion of key extracellular matrix (ECM) components in cells harboring the SWS mutation (Xia et al., 2022). However, this system had inherent limitations. As a transformed and terminally differentiated cell line, SW1353 cells underwent early apoptosis upon spheroid formation, limiting their capacity to model sustained chondrogenesis and matrix maturation. Consequently, this approach did not fully capture the dynamic processes underlying cartilage development.

Animal models have similarly provided incomplete insight into the human skeletal phenotype. Although patients with heterozygous *COG4* mutations develop skeletal dysplasia, here, we generated *Cog4^+/G512R^* and *Cog4^G512R/G512R^* mouse lines, in which heterozygous animals do not exhibit skeletal abnormalities, while homozygous animals display embryonic lethality accompanied by severe skeletal defects including shortened bones, reduced mineralization, and impaired endochondral ossification. These findings suggest species-specific differences in gene dosage sensitivity and highlight the limitations of murine models in recapitulating the human disease. This motivated us to establish patient-derived iPSC cartilage organoids to study disease mechanisms in a human developmental context.

Several in vitro strategies have been developed over the past decade to differentiate iPSCs into chondrocytes, both to model skeletal development and disease and to support regenerative medicine applications (C. L. Wu et al., 2021; Guzzo & Drissi, 2015; Dicks et al., 2023; Khan et al., 2023; Lamandé et al., 2023; Zujur et al., 2023). Here, we generated chondrocytes from iPSCs via mesenchymal progenitors followed by high-density 3D pellet culture, an approach that closely mimics mesenchymal condensation and promotes cell–cell interactions required for chondrogenesis (Caron et al., 2012; Grigull et al., 2020; Guzzo et al., 2013; Koyama et al., 2012; Ullah et al., 2012). This approach demonstrated progressive induction of canonical chondrogenic markers and robust extracellular matrix deposition in control organoids, consistent with established chondrocyte differentiation trajectories and late appearance of hypertrophic chondrocytes that facilitate endochondral ossification.

Recent work by Lawrence et al. generated a single-cell RNA sequencing atlas of embryonic long bones and integrated these data with in vitro chondrogenesis datasets, providing a comprehensive reference for endochondral ossification (Lawrence et al., 2025). Consistent with embryonic stem cell-derived and bone marrow MSC differentiation datasets (Huynh et al., 2019; Lawrence et al., 2025), our control cultures displayed progressive induction of chondrogenic genes including *SOX9*, *COL9A1*, *COL2A1*, *ACAN*, *MATN3*, and *COMP*. However, unlike datasets that progress to hypertrophic or osteogenic stages, our cultures did not show robust activation of hypertrophic transcription factors (*RUNX2*, *DLX5*, *SP7*) or osteoblast markers (*ALPL*, *BGLAP*, *IBSP*). Instead, we observed expression of pre-hypertrophic markers such as *COL10A1*, *PTH1R*, and *SPP1*, suggesting progression to early hypertrophic stages without full osteogenic commitment. Similar differentiation dynamics have been reported in single-cell analyses of iPSC-derived chondrocytes, where cells preferentially occupy proliferative, resting, or articular-like states rather than rapidly transitioning to hypertrophy (Wu et al., 2021b). These results indicate that our system models early to mid-stages of endochondral differentiation rather than terminal ossification, aligning with articular or growth plate-like trajectories.

In contrast to control lines, SWS organoids exhibited marked defects in activation of skeletal development programs and NABA core matrisome genes. Temporal expression patterns suggest that differentiation is initiated but fails to progress, resulting in a developmental block. Gene expression profiles in patient samples at Day 28, including persistent expression of *THY1, COL1A1, COL1A2, COL3A1, SPARC, POSTN, BMP4,* and *FGFR2*, are consistent with retention of mesenchymal stromal/progenitor characteristics rather than terminal chondrocyte identity (Huynh et al., 2019; Katsumata et al., 2017; Muntión et al., 2024). Consistent with these findings, CODEX analysis demonstrated reduced numbers of SOX9, ACAN, and COL2A1 positive cells together with persistence of mesenchymal-associated markers such as CD9, CD44, and COL1A1. Collectively, these transcriptional and proteomic signatures indicate that SWS organoids remain in a mesenchymal progenitor-like state rather than progressing toward mature chondrocytes.

To connect these differentiation defects to the underlying *COG4* variant, we examined glycosylation changes in this system using a novel spatial multiomic approach by simultaneously capturing protein and glycosylation profiles at sub-cellular resolution. While earlier SWS models reported subtle N-glycosylation alterations, lectin profiling in our system revealed broad changes across a range of detectable glycosylation moieties, including reductions in complex glycan motifs and increased high-mannose structures. The most striking change, however, involved decorin, consistent with prior SWS studies (C. R. Ferreira et al., 2018; Xia et al., 2021, 2022). Disaccharide analysis of chondroitin sulfate proteoglycans (CSPGs) revealed an approximately 90% reduction in total glycosaminoglycan (GAG) chains in SWS organoids, indicating markedly impaired GAG modification of decorin and other CSPGs.

Because CSPGs regulate growth factor availability and cell-matrix interactions essential for cartilage formation, loss of GAG chains is expected to disrupt signaling environments and matrix assembly required for robust chondrocyte differentiation (Hildebrand et al., 1994; Yamaguchi et al., 1990; Schwartz & Domowicz, 2022). Consistent with this interpretation, variants affecting GAG linker synthesis, polymerization, or sulfation are well known causes of skeletal dysplasias characterized by short stature and abnormal cartilage development (Dubail & Cormier-Daire, 2021; Paganini et al., 2019, 2020). Decorin and related CSPGs are critical for collagen fibrillogenesis and matrix stabilization, and reduced GAG modification leads to diminished aggrecan retention and compromised biomechanical properties (Chen et al., 2020; Reed & Iozzo, 2002; Reese et al., 2013). Collectively, our data suggest that insufficient GAG modification of decorin and other CSPGs contributes directly to defective ECM assembly and impaired chondrogenesis in SWS organoids, providing a plausible mechanism linking COG4 dysfunction to skeletal abnormalities (Figure 7).

**Figure 7:**
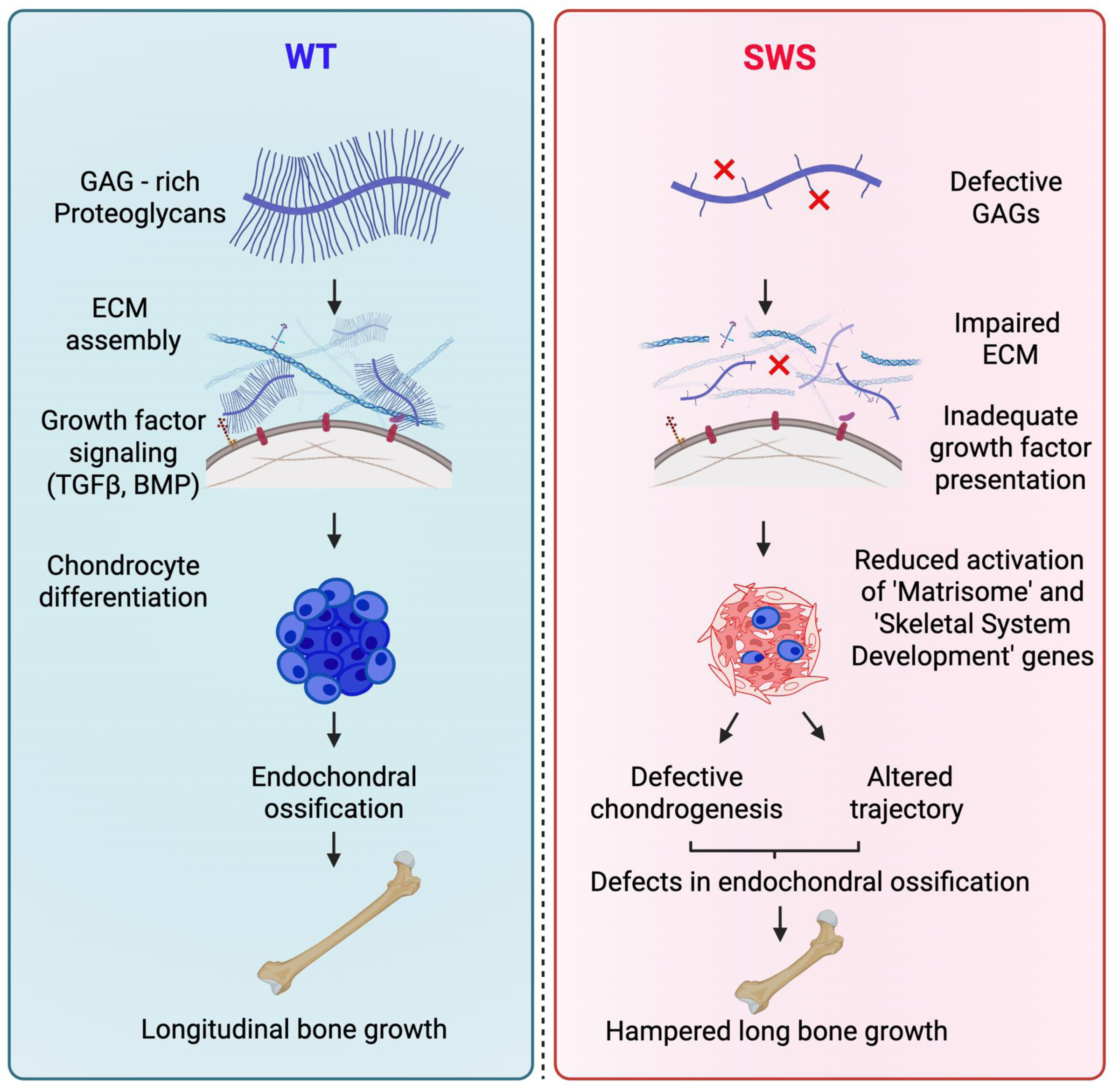
Schematic showing the proposed model for disease pathogenesis.

These findings suggest that disruption of Golgi-dependent ECM maturation imposes a specific developmental constraint during chondrogenesis. In this context, it is informative to compare SWS with other skeletal disorders that affect distinct stages of the chondrogenic trajectory.

Notably, SWS organoids exhibit differentiation arrest at very early stage during chondrogenic commitment. This arrest coincides with the stage at which control cultures undergo rapid ECM expansion, suggesting that chondrocytes encounter a secretory burden requiring high Golgi processing capacity. This phenotype contrasts with other well-characterized disorders. For example, activating mutations in FGFR3, as seen in achondroplasia, permit chondrogenic differentiation but restrict proliferative expansion of growth plate chondrocytes through aberrant MAPK signaling (Ornitz & Legeai-Mallet, 2017). In contrast, several skeletal disorders including type II collagenopathies and pseudoachondroplasia caused by COMP mutations is characterized by ER stress due to intracellular retention of matrix proteins and disrupted ECM assembly despite relatively preserved early differentiation (Rellmann & Dreier, 2018). Other disorders, including TRPV4-associated skeletal dysplasias, primarily affect later stages of chondrocyte hypertrophy and terminal differentiation (Kang et al., 2012). In contrast, conditions such as hypophosphatasia impair matrix mineralization downstream of chondrogenesis (Whyte, 2016).

More broadly, these findings place SWS within a growing class of disorders in which defects in ER–Golgi trafficking disproportionately affect tissues with high secretory and ECM demands, such as cartilage and bone (Tüysüz, 2025). Our data therefore suggest that efficient Golgi-dependent proteoglycan maturation represents a critical checkpoint during human chondrogenesis. Disruption of this process likely prevents progression from early chondrogenesis to a matrix-rich, growth plate-like state, thereby contributing to skeletal dysplasia despite apparently normal early lineage specification.

Overall, our results demonstrate that patient-specific iPSC-derived cartilage organoids recapitulate disease-relevant and species-specific phenotypes not captured by existing cell line or murine models. Beyond confirming defective decorin glycosylation, our findings identify a previously unrecognized link between Golgi-dependent proteoglycan maturation and a developmental checkpoint in human chondrogenesis, with direct implications for skeletal disease pathogenesis. More broadly, this work highlights the importance of studying disease mechanisms in lineage-relevant human models. Finally, the integration of iPSC-based chondrogenesis, multiplexed antibody and lectin profiling provides a versatile platform for investigating skeletal disorders driven by extracellular matrix and glycosylation defects.

### Limitations of the study

While our study provides mechanistic insight into altered proteoglycan glycosylation and extracellular matrix defects using patient-derived cartilage organoids, several limitations should be considered. First, the analysis was performed using a limited number of patient-derived iPSC lines, reflecting the rarity of Saul-Wilson syndrome, and may not fully capture the spectrum of phenotypic variability. Second, although the organoid system models human chondrogenesis in a controlled developmental context, it does not fully recapitulate the complexity of in vivo cartilage. Our analyses were focused on Day 14 as an early time point, at which control and patient organoids are comparable in size, although molecular differences begin to emerge at or before this stage and become more pronounced at later time points.

Earlier time points would further refine the temporal onset of these defects. Within this framework, our data support defects in decorin glycosylation and chondroitin sulfate modification as contributors to impaired matrix assembly and chondrocyte commitment. While we did not directly test whether restoration of proteoglycan glycosylation can rescue these phenotypes, this represents an important direction for future studies. Despite these limitations, our model provides a human, disease-relevant platform to interrogate early cartilage defects and establishes a foundation for future mechanistic and therapeutic studies.

## Resource Availability

### Lead Contact

Further information and requests for reagents may be directed to the Lead Contact, Hudson Freeze.

### Materials Availability

Cell lines generated and used in this study are available upon reasonable request from the Lead Contact.

### Data and Code Availability

The RNA sequencing data generated in this paper is uploaded to GEO with accession number: GSE306715.

## Acknowledgements

The authors would like to thank the NIH funding R01DK99551 and The Rocket Fund, to support this work. The research was also supported in part by the Intramural Research Program of the National Institutes of Health (ZIA HD009024 to C.R.F.). The contributions of the NIH author(s) are considered Works of the United States Government. The findings and conclusions presented in this paper are those of the author(s) and do not necessarily reflect the views of the NIH or the U.S. Department of Health and Human Services. We thank the core facilities– Guillermina Garcia (Histology Core, SBP), Yoav Altman (Flow Cytometry Core, SBP), Biswa Choudhury and Mousumi Paulchakrabarti (GlycoCore, UCSD) for their dedicated services. The authors thank Dr. Evan Snyder, Dr. Anne Bang, and Dr. Yu Yamaguchi for critical discussion. We also thank Jamie Smolin for excellent technical assistance.

## Authors contributions

S.M. and H.H.F. conceptualized the project, designed the experiments, analyzed the data and oversaw all studies.

S.M. performed most of the experiments.

S.M. and S.A. conducted CODEX optimization and experiments, W.W. oversaw CODEX experiments and data analysis.

J.T., R.K., and G.A. performed the mouse experiments, and C.R.F. conceptualized and oversaw mouse generation and data analysis for these experiments.

R.M. analyzed the RNA-seq data and prepared the corresponding figures.

S.M. wrote the manuscript. All authors reviewed and contributed to editing the manuscript.

## Declaration of interests

The authors have no competing interests to declare.

## Declaration of generative AI and AI-assisted technology in the writing process

During the preparation of this manuscript, the authors used ChatGPT-5 mini from OpenAI for language correction and to tighten some sections to meet the *Cell Stem Cell* article word limit. After using this tool, the authors reviewed and edited the content as needed and take full responsibility for the content of the publication.

## STAR★Methods

### Experimental models and study participant details

#### iPSC Cultures

Two patient and two control fibroblasts were reprogrammed to iPSCs in Dr. Evan Snyder’s lab. The undifferentiated hiPSCs were cultured in mTeSR1™ plus medium (Stem Cell™ Technologies, 100-0276) in 6 cm petri dishes coated with 0.28 mg/ml Geltrex™ Matrix solution (Gibco A1413202) at 37 °C in a 5 % CO_2_ atmosphere. The medium was changed every day and cells were sub-cultured every 4–5 days using ReLeSR™(Stem Cell™ Technologies, 100-0483).

#### Differentiation of iPSCs to MPCs

Mesenchymal progenitor cells (MPC) were generated from iPSCs using the STEMdiff Mesenchymal Progenitor kit (Stem Cell Technologies, 05240) in accordance with the manufacturer’s instructions. In brief, induced pluripotent stem cells (iPSCs) were cultured in mTeSR1 media (StemCell Technologies) until they reached approximately 40–50% confluency. iPSCs were then incubated in STEMdiff-ACF mesenchymal induction medium for 3 days, followed by an additional 3 days in MesenCult-ACF Medium. On day 6, early mesodermal progenitor cells were plated in wells coated with MesenCult-ACF Attachment Substrate and maintained in MesenCult-ACF Medium. Cells were split at 80% confluency and characterized by flow cytometry 20–22 days post-differentiation using Human Mesenchymal

Stem Cell Multi-Color Flow Cytometry Kit (R&D Systems, FMC002). Single-cell suspensions of MPC were stained with a panel of immunophenotyping antibodies including Mesenchymal cell positive markers (CD146/MCAM, CD90/Thy1 and CD73) and Mesenchymal cell negative marker (CD45) and pluripotency marker (Oct 3/4). Briefly, 1 x 10^6^ cells were stained with 1ug of each antibody for 30-45 minutes at room temperature in the dark. Following the incubation, cells were washed and resuspended in 250 μL of Staining Buffer for flow cytometric analysis.

#### Differentiation of MPCs to 3D cartilage organoids

For chondrogenic differentiation, a total of 5 × 10^5^ MPCs were resuspended in 0.5 ml of MesenCult™-ACF Chondrogenic Differentiation Medium (Stem Cell Technologies, 05455) and centrifuged into a 15 mL polypropylene tube as a cell pellet and incubated at 37°C and 5% CO_2._ On Day 3, gently add 0.5 ml of fresh medium to the pellets, for a final volume of 1ml. On Day 6 and every 3 days afterwards, the medium was gently aspirated and replaced with 0.5 mL of medium until Day 35. At selected time points, the chondrogenic organoids were collected and processed based on downstream assay.

### Method details

#### BFA-induced retrograde transport assay using Imaging Flow Cytometry

MPCs were detached from the plate using ACF Enzymatic Dissociation Solution and washed with prewarmed growth medium. Cells were then incubated with growth medium containing 0.25 μg/ml BFA for 0, 15, 22, 30 and 45 min at 23°C. The incubations were stopped by washing cells with DPBS and fixation was done with 4% paraformaldehyde for 10 min at room temperature. Cells were permeabilized with 0.4 % Triton X 100 for 10 minutes and stained by immunofluorescence using Alexa Fluor 488 anti-Giantin antibody (BioLegend) and DAPI for DNA staining. Cells were imaged using multispectral imaging flow cytometry (ImageStreamX mark II imaging flow-cytometer; Amnis Corp, Seattle, WA).

Approximately 1 × 10^4 cells were collected from each sample, and data analysis was performed using image analysis software (IDEAS 6.2; Amnis Corp), following the methodology outlined in Wortzel et al. 2017. In brief, firstly the compensation was done for fluorescent dye overlap using single-stain controls. Cells with a positive DAPI signal were gated for single cells, using the area and aspect ratio features, and for focused cells, using the Gradient RMS feature. To quantify Golgi fragmentation, two specific features were selected based on Giantin staining: Minor Axis Intensity (the intensity-weighted narrowest dimension of the best-fit ellipse) and Area (the number of microns squared within a mask). These metrics were calculated using the Threshold_60 mask, which encompassed the 60% highest intensity pixels of the Giantin staining. The two features were then plotted on a bivariate plot, and different Golgi morphologies (Intact, Partial, and Diffuse) were gated through visual inspection (Figure S3C).

#### mRNA Expression Analysis

Total RNA was extracted from cartilage organoids using TRIzol^TM^ (ThermoFisher, 15596081) reagent according to manufacturer’s protocol. cDNA was synthesized using SuperScript IV VILO (SSIV VILO) Master Mix (Invitrogen™, 11766050). qPCR was performed using TaqMan assay probes and TaqMan™ Universal PCR Master Mix (Applied Biosystems™, 4304437).

The mRNA levels were normalized to the levels of housekeeping gene, GAPDH, and 2^−ΔΔCt^ values were calculated and compared.

#### RNA sequencing library preparation

RNA sequencing was performed at iMPC stage and two different time points of chondrogenic differentiation (Day 14 and Day 28) to map the transcriptional changes in control vs patient samples. Total RNA was extracted from the cartilage organoids using TRIzol™ (ThermoFisher, 15596081) reagent according to manufacturer’s instructions. Standard mRNA library was prepared using the PolyA selection and the data was generated at the UC San Diego IGM Genomics Center utilizing an Illumina NovaSeq X Plus that was purchased with funding from a National Institutes of Health SIG grant (#S10 OD026929). Bioinformatic analysis was performed in the Sanford Burnham Prebys Bioinformatics Core.

#### RNA-seq analysis

RNA-seq samples were processed using nf-core rnaseq v3.14.0 [PMID: 32055031] Nextflow pipeline with parameters "--aligner star_rsem --igenomes_ignore --genome null". Human genome version GRCh38 (primary assembly) and Gencode version 45 (primary assembly) annotations were used with nf-core rnaseq pipeline. Singularity software containers were used for nf-core pipeline run using “-profile singularity” parameter. Differential gene expression analysis was performed using nf-core differentialabundance pipeline version v1.5.0 (Ewels et al., 2020) with parameters ‘--filtering_min_abundance 4, --filtering_min_proportion 0.50, and --differential_min_fold_change 2.0”. Genes with a minimum fold-change of 2.0 and FDR < 0.05 were considered differentially expressed.

Time-course gene expression analysis was performed using maSigPro v1.76.0 in R v4.4.1 (Nueda et al., 2014). Genes with RNA-seq counts equal or more than 10 counts in at least 12 samples (50% of all samples) were retained for maSigPro analysis. MPC samples were used as initial time point (time = 0) for the analysis. Time-course analysis was performed using maSigPro “p.vector(counts, design_matrix, counts=TRUE, Q = 0.05)” and “T.fit” functions.

#### Immunohistochemistry

The chondrogenic pellet was fixed with 4% formaldehyde, followed by subsequent standard paraffin embedding. Alcian blue staining was performed by Histology core at Sanford Burnham Prebys (SBP) following standard protocol.

#### CODEX staining and data analysis

Multiplex CODEX imaging was performed on fresh-frozen OCT-embedded tissue sections of control and patient-derived organoids. Organoids were harvested at defined time points and fixed overnight in 0.5% formaldehyde. Following washes in phosphate-buffered saline (PBS), samples were treated with 30% (w/v) sucrose in PBS. The tissues were then embedded in two 25 x 25 mm molds filled with optimal cutting temperature (OCT) compound and rapidly frozen in a bath of semi-frozen isopentane cooled in liquid nitrogen. Cryosections were collected onto gelatin-coated CODEX coverslips and processed for staining according to the manufacturer’s protocol (Akoya Biosciences), with minor modifications to the staining steps. A customized CODEX antibody panel targeting cartilage markers, along with a panel of glycan-binding lectins (GLYPH), each conjugated to unique DNA barcodes, was used for multiplexed imaging. Acquired images were processed using the CODEX post-processing pipeline. Cell segmentation was performed using an optimized script to generate CRISP outputs from extended depth-of-field images derived from deconvolved optical slices after background subtraction to remove autofluorescence. Downstream analyses were conducted using custom scripts implemented in Jupyter notebooks.

#### Immunoblotting

Conditioned medium from cartilage organoids was collected, concentrated, and samples were prepared with 5X SDS buffer. Equal amounts of proteins were separated via SDS–polyacrylamide gel electrophoresis followed by transfer and antibody inoculation. Antibodies used were: Decorin (R&D, MAB143) and Aggrecan (Biolegend, 674902).

#### Generation of *Cog4^+/G512R^* mice

*Cog4^em1Cferr^* (abbreviated *Cog4^+/G512R^*) knock-in mice modeling the human *COG4* c. c.1546G>C (p.G516R; NM_015386.2; rs1555575860) variant were generated using CRISR-Cas9-mediated homology-directed repair (HDR). A mix of guide RNA (5’-CGGCACTTGTCACCCCACGC-3’, synthesized as IDT Alt-R RNA Oligo), a donor DNA repair template (0.25 µM each), and spCas9 was used for electroporation into FVB/N × C57BL/6J F1 hybrid eggs. The repair template was synthesized as IDT Ultramer single-stranded DNA fragments, with 84 bp and 108 bp isogenic homology arms flanking the two nucleotide changes introduced by the donor repair template. The full reverse complement sequence of the 200-bp repair template is: 5’-GTCTGCGTCTGCCACTCTGCCCCACAGGGATGTTCTATGTAATAAGCTAAGAATGGGCTTCCCAGCCACCACCTTACAGGACATtCAGCGTcGGGTGACAAGTGCCGTGAACATCATGCACAGCAGCCTCCAGCAGGGCAAGTTTGACACCAAAGGCATCGAGAGCACTGATGAAGCCAAGCTGTCCTTCCTGGTAAGTG-3′ (lower case indicates the two nucleotide changes compared to wild-type). The resulting allele, *Cog4^em1Cferr^*, also referred to as *Cog4^+/G512R^*, contains two nucleotides different from wild-type (GRCm38 chr8:110868259-110868266cCAGCGTg>tCAGCGTc) resulting in a single amino acid change in exon 12 of mouse COG4 p.Gly512Arg, equivalent to human COG4 p.Gly516Arg.

Sanger sequencing was used to confirm the knocked-in allele in founder animals (forward primer: 5’-ggtgTTGAACAGTCTGCGTC-3’; reverse primer: 5’-CCCTTGAGAAAATCTGCCCC-3’). Indels in the founder animals were ruled out by fluorescent PCR (forward primer with M13F tail: 5’-TGTAAAACGACGGCCAGTggtgTTGAACAGTCTGCGTC-3’; reverse primer with PIG tail: 5’-GTGTCTTCCCTTGAGAAAATCTGCCCC-3’; universal M13F-FAM primer: 5’-FAM-TGTAAAACGACGGCCAGT-3’) (Carrington et al., 2015). For genotyping experimental animals, DNA was extracted from pup tail biopsies and DNA prepared using a rapid DNA extraction kit (REDExtract-N-Amp Tissue PCR Kit; Sigma-Aldrich, MO, USA). Samples were amplified using a TaqMan Fast Advanced Master Mix for qPCR (ThermoFisher Scientific, MA, USA) with a VIC-labeled probe for wild-type sequence and a FAM-labeled probe for alternate sequence (ThermoFisher Assay ID: ANFV2GU). An ABI StepOne Plus instrument was used for thermocycling and detection.

#### Study approval

Mice were maintained in a specific pathogen-free AAALAC-accredited facility, and all procedures were approved by the Institutional Animal Care and Use Committee (ACUC) of the National Human Genome Research Institute (NHGRI) or the National Institute of Dental and Craniofacial Research (NIDCR).

#### Morphometric measurements

Weights were measured using a Sartorius BP3100S balance (readability: 0.01 g), and lengths were measured using Reed R7400 digital caliper (accuracy: 0.03 mm).

#### Dual-energy X-ray absorptiometry (DXA)

An UltraFocus DXA cabinet (Faxitron®, AZ, USA) was used to measure whole body bone mineral density (BMD) of WT and *Cog4^+/G512R^* mutant mice. Following the calibration procedure as per the manufacturer’s manual, X-ray scans were imaged at 4X magnification.

#### MicroCT analysis of mouse bones

Femur bones from *Cog4^+/G512R^* female mice and littermate controls (24-28 weeks) were dissected and fixed with Z-fix fixative (Anatech, MI, USA) for 24 h, then rinsed twice with 1X PBS, pH 7.4 (Thermo Fisher Scientific) and stored in 70% ethanol at 4°C. Scans were obtained with a SCANCO µCT 50 machine and ex vivo microCT analyses were performed with the BMA add-on module of the Analyze 14.0 Software (AnalyzeDirect, KS, USA). The region of interest (ROI) was set using a manually-determined global threshold. Three-dimensional microstructural bone properties were calculated according to the manufacturer’s software. Statistical analysis was performed using an unpaired Student’s t-test and p values < 0.05 were considered statistically significant.

#### Whole-mount skeletal staining

Fresh staining solutions of 0.005% (w/v) Alizarin red in 1% KOH, 0.03% (w/v) Alcian blue 8GX in 80% ethanol/20% glacial acetic acid, and 1% KOH in deionized H_2_O were prepared.

Embryos at E15.5 were collected and placed in vials with 1x PBS and briefly immersed in 70°C water bath before removal of tissue. Defleshing of the carcasses was manually performed, removing as much soft tissue as possible, before fixing the embryos in 95% ethanol at room temperature overnight. The embryos were transferred to 100% acetone and incubated at room temperature for a second night, before rinsing the embryos with deionized H_2_O and stained with enough Alcian blue stain to cover the body, staining for a third night. The embryos were washed twice in 70% ethanol for an hour each, then incubated in 95% ethanol overnight. The ethanol was removed, and 1% KOH was added for an hour until the tissues were visibly cleared at room temperature. After removing the 1% KOH, the embryos were counterstained with Alizarin red solution for four hours, after which the embryos were stored in 50% glycerol in 1% KOH solution at room temperature overnight. The embryos were transferred to 50% glycerol in ethanol storage before photodocumentation. Skeletal preparations were positioned in lateral orientation and photographed for morphometric measurements. The images were captured using a digital camera mounted on a Zeiss SteREO Discovery.V12 microscope (Carl Zeiss AG, Oberkochen, Germany).

#### Embryo collection for CODEX

The embryos representing all genotypes: wild-type (n = 3), heterozygous (n = 7) and homozygous (n = 3) were collected at embryonic day 15.5 (E15.5). After removal of the skin and internal organs, the skeletons were fixed in 0.5% paraformaldehyde (PFA) in PBS for 24 hours at room temperature (RT). Samples were rinsed twice with PBS and subsequently decalcified with 0.5M EDTA on a shaker at RT for 3 days. Following decalcification, EDTA solution was replaced by 30% (w/v) sucrose in PBS and samples were stored at 4°C until all the tissues have sunken to the bottom of the tube. The tissues were then embedded in OCT compound, as described for organoids.

## Notes

### Competing Interest Statement

The authors have declared no competing interest.

