## Supplementary material for "Patient iPSC-Derived Cartilage Organoids Reveal Defective ECM Deposition and Altered Chondrogenic Trajectory in Saul-Wilson Syndrome": Figure S1 - S6

5, Co-senior author

6, Lead contact

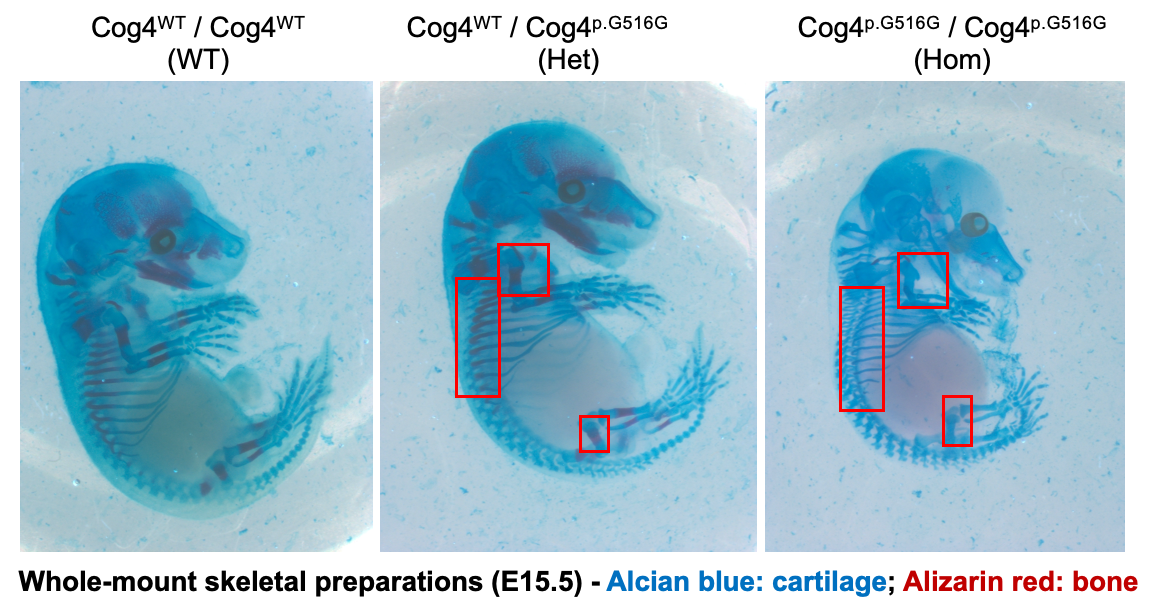

B

A

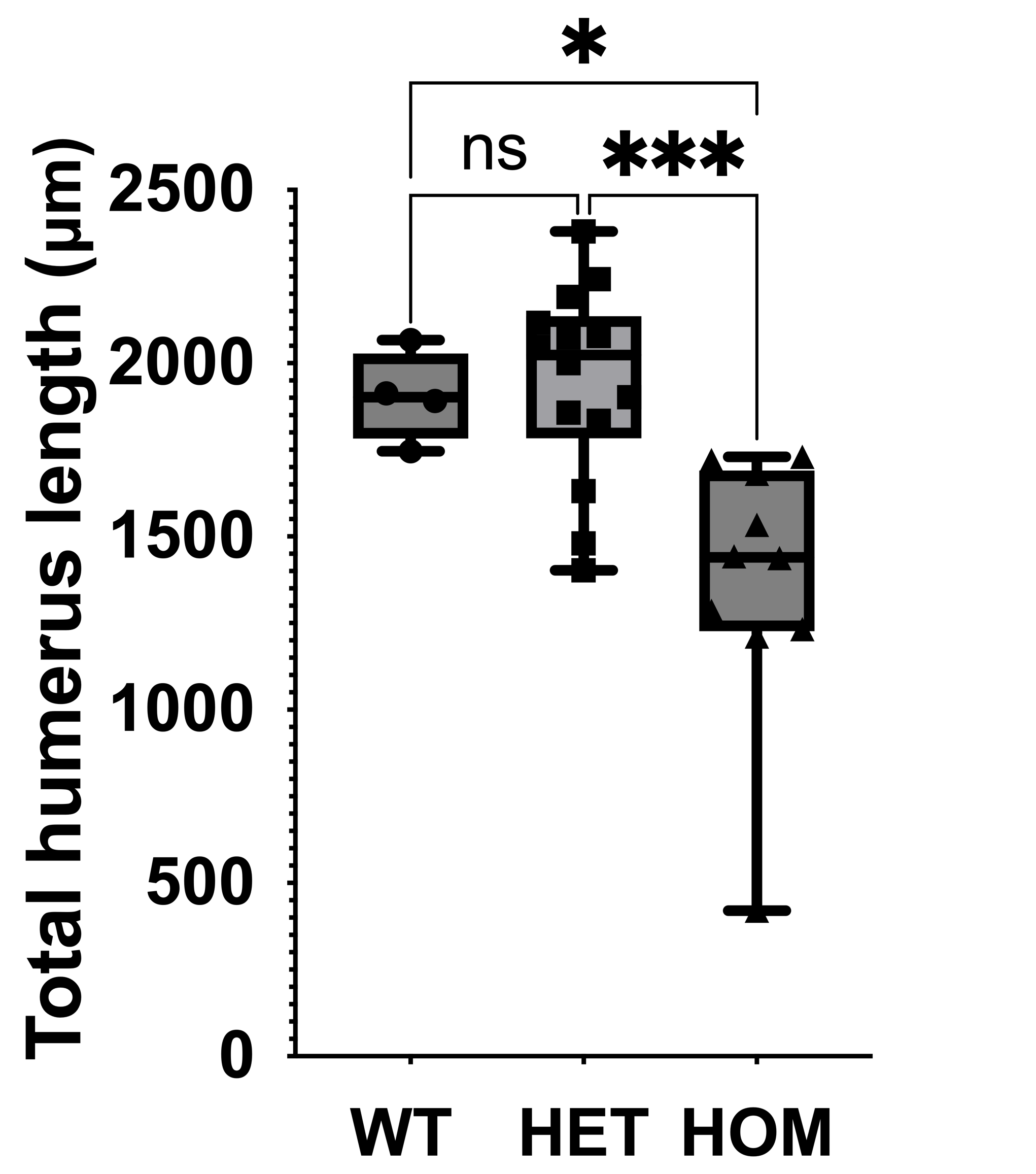

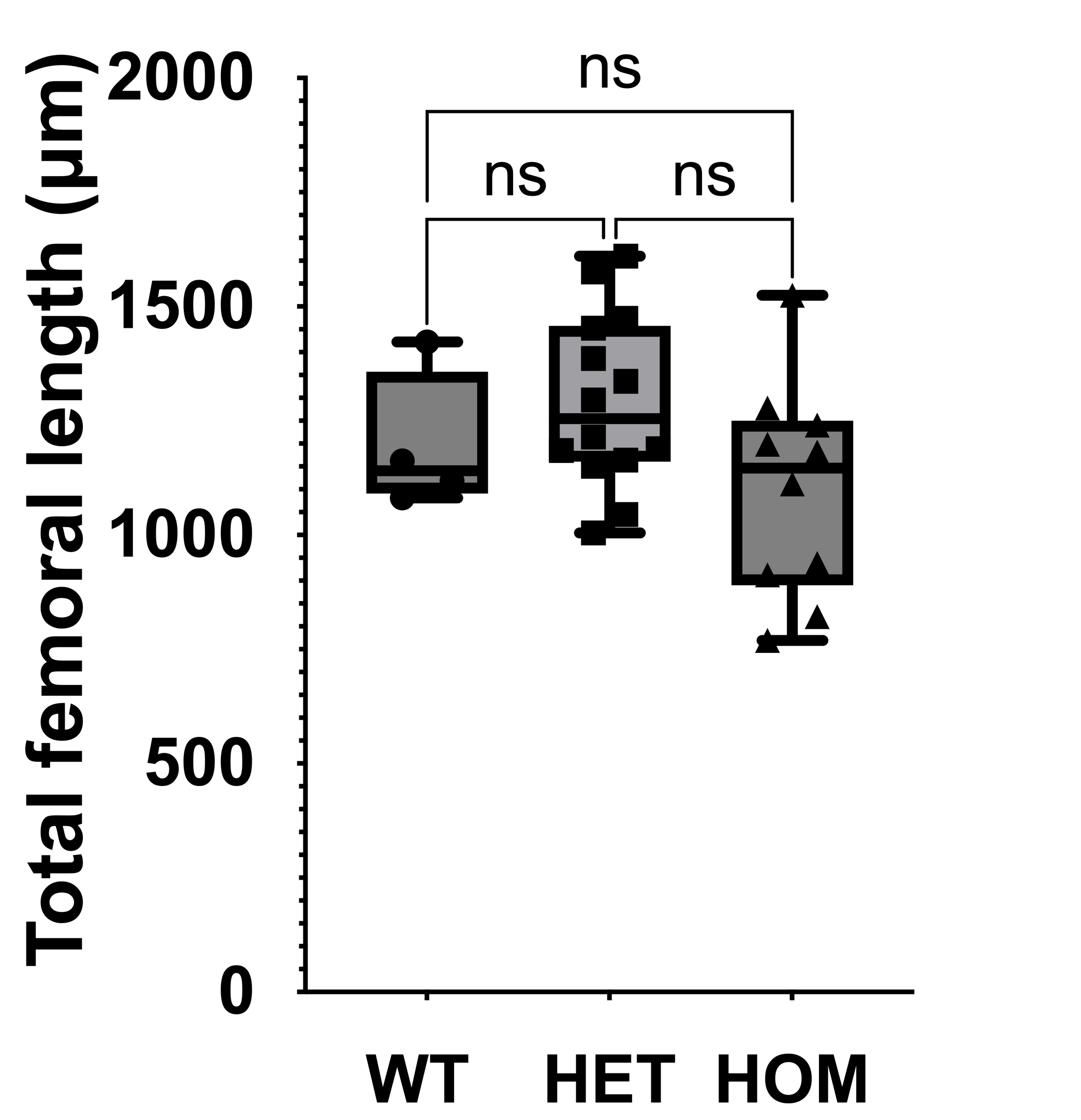

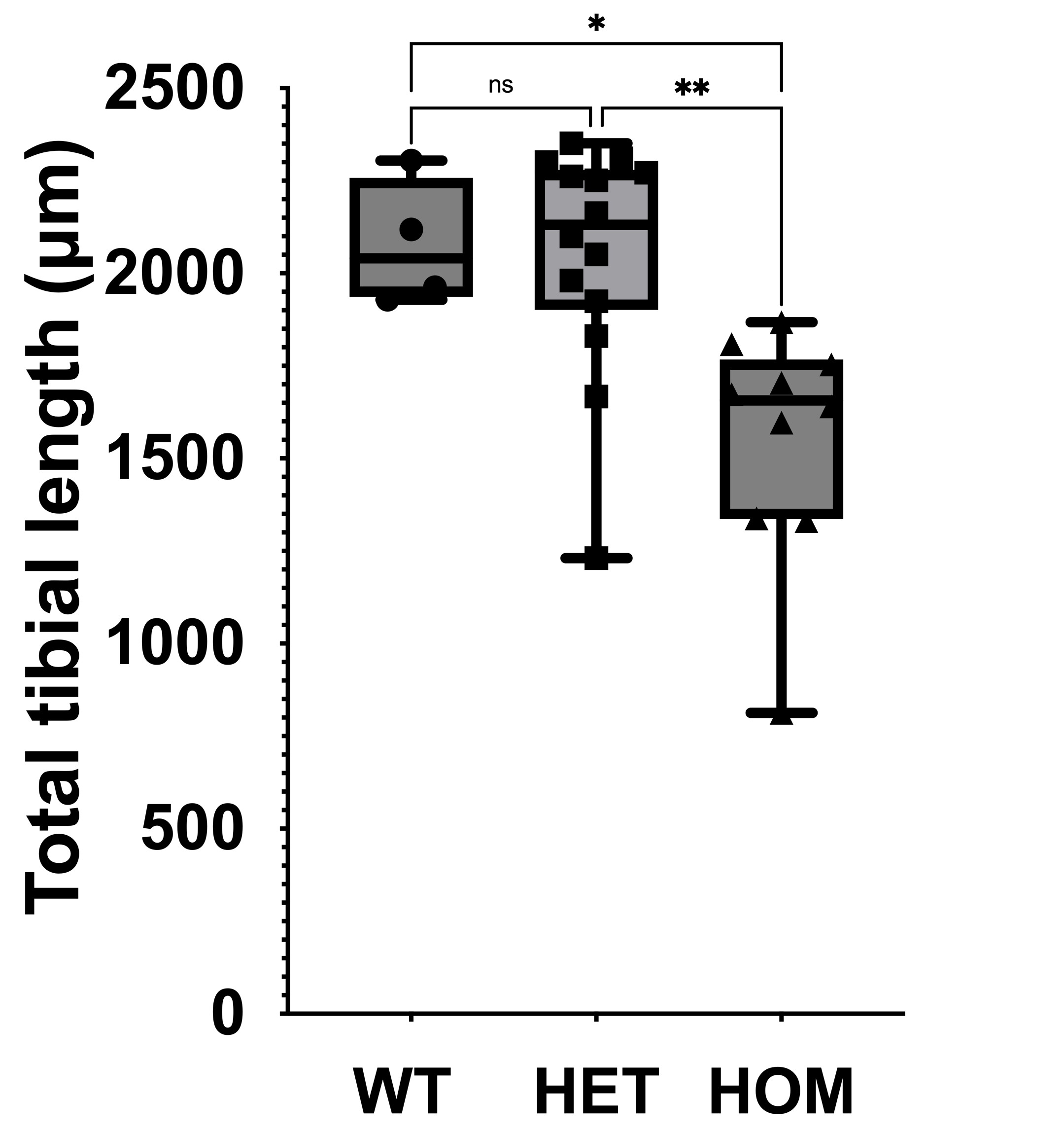

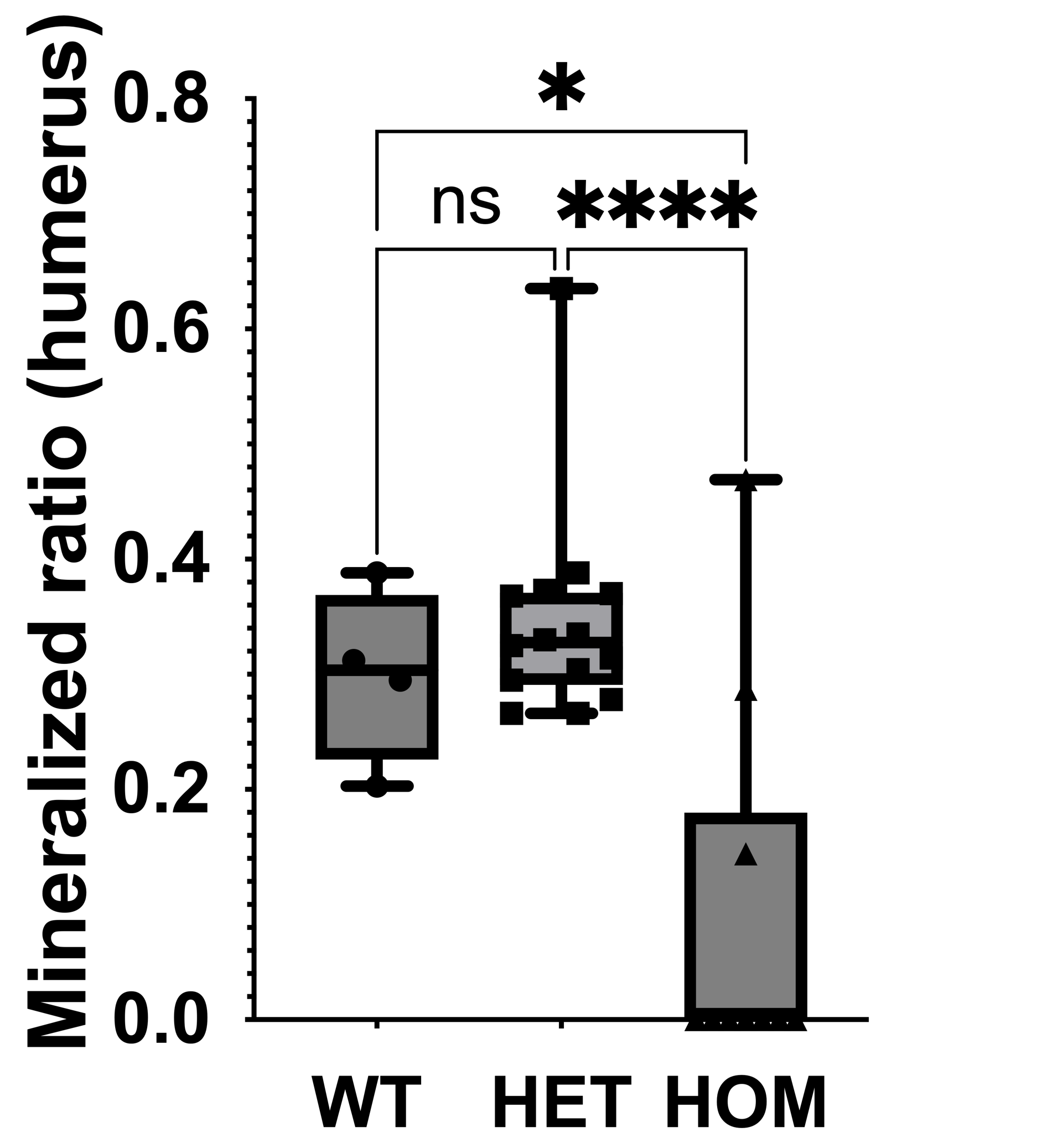

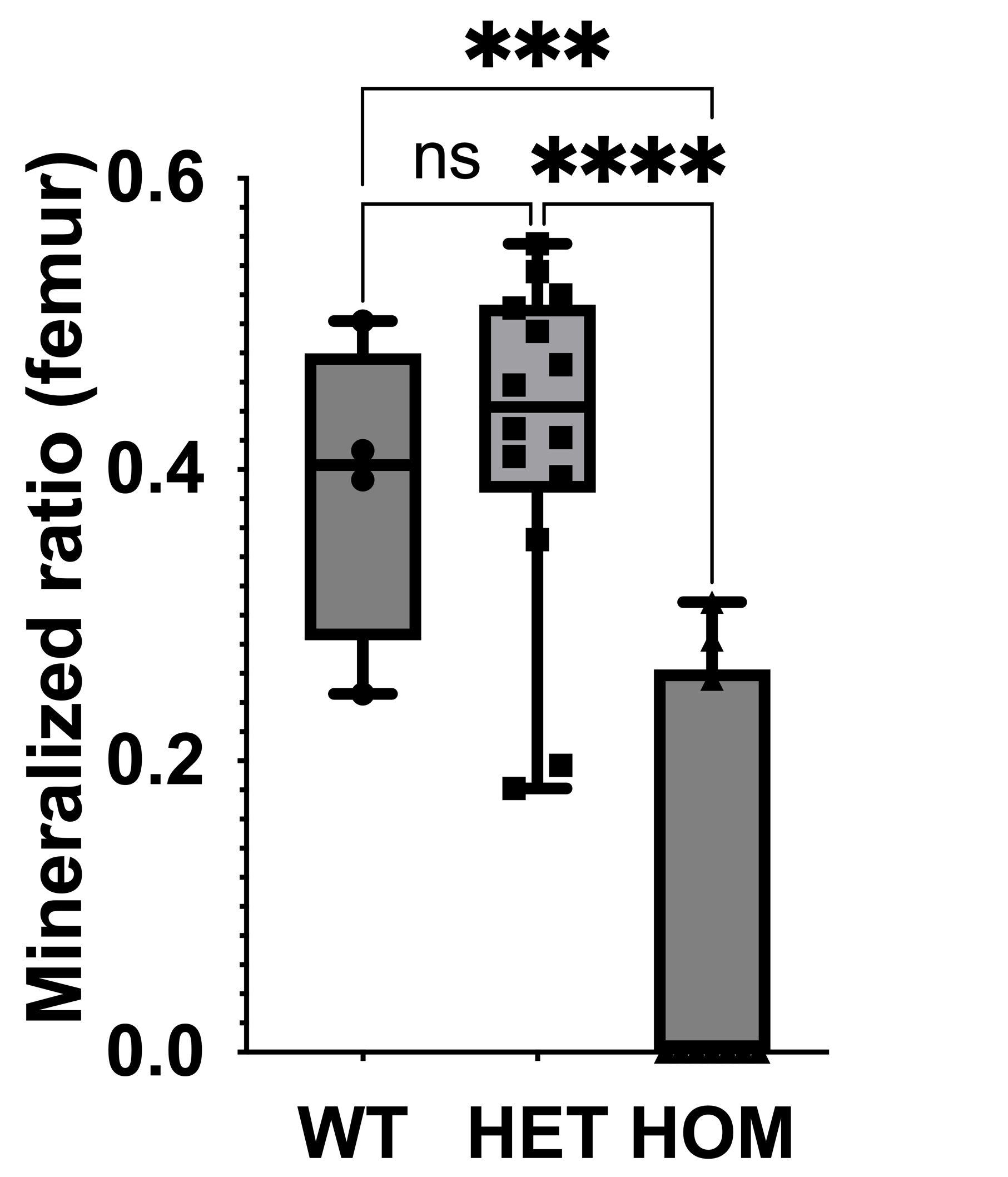

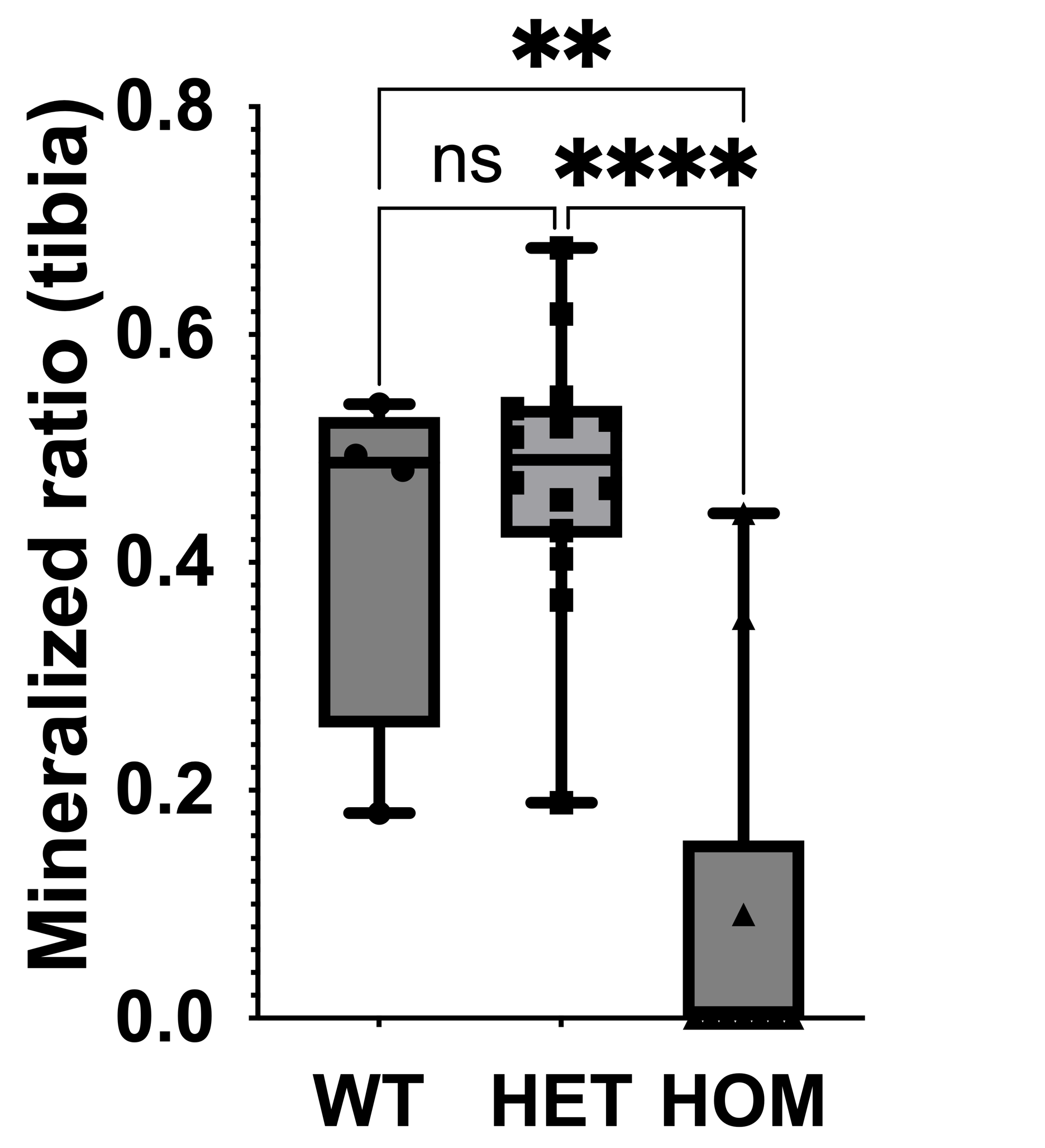

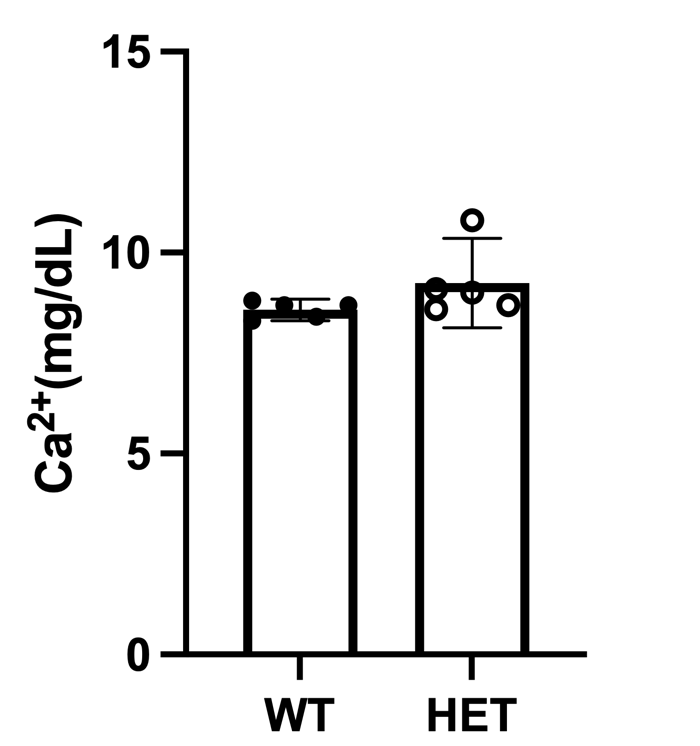

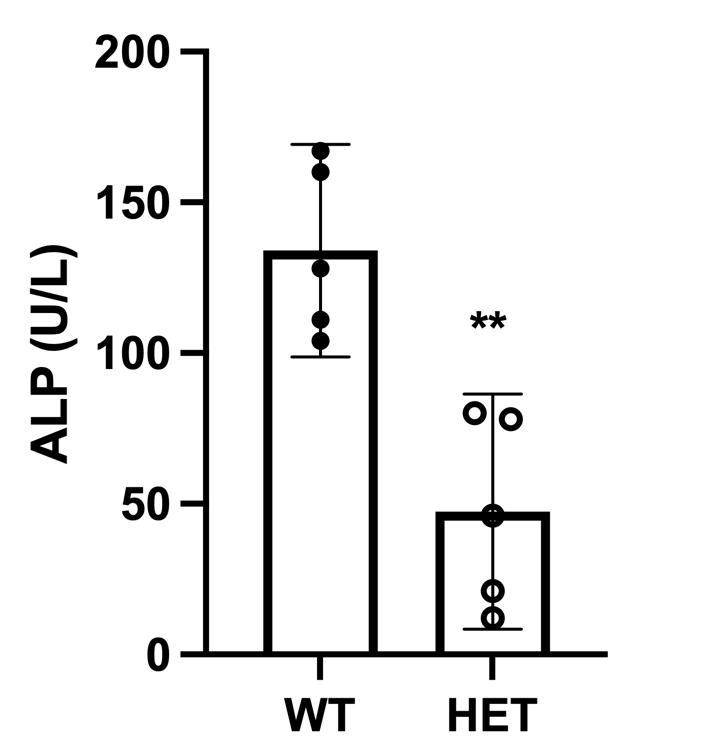

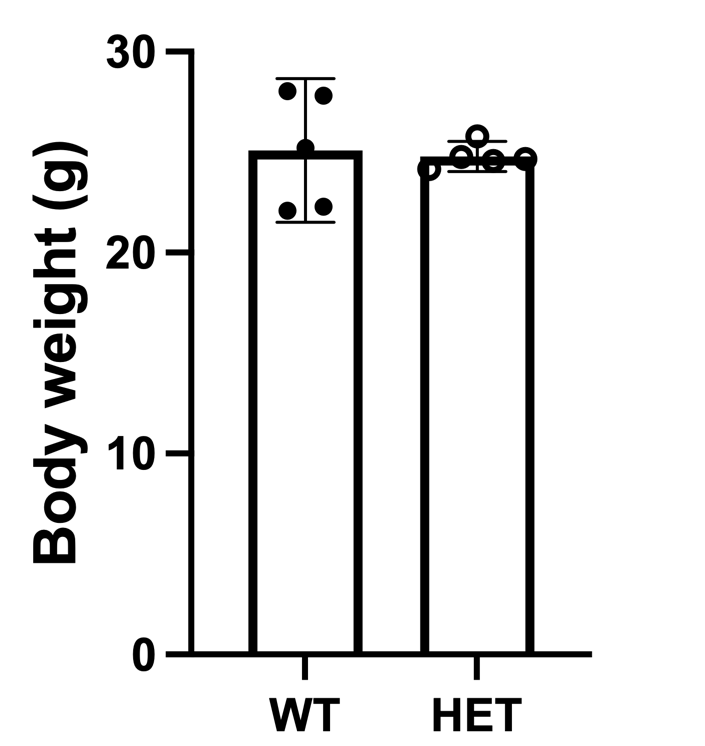

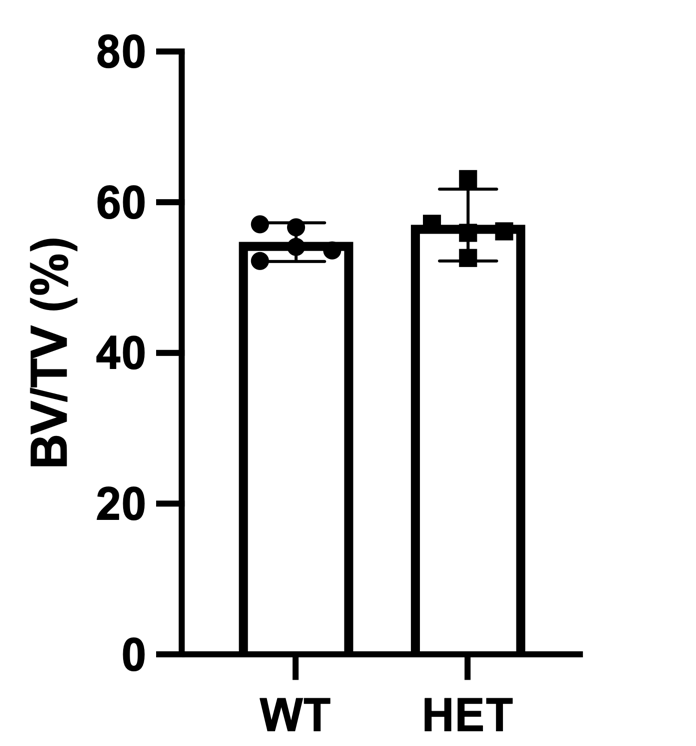

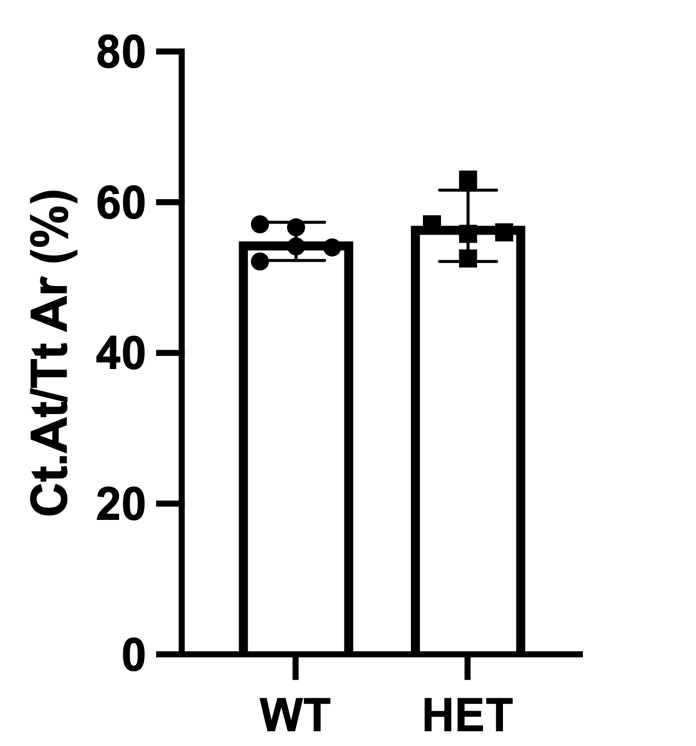

C

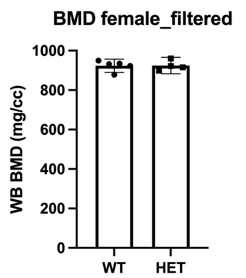

**Figure S1**

A. Whole-mount skeletal preparations of E15.5 embryos. Cartilage is stained blue with Alcian blue, while the mineralized bone is stained red with Alizarin red.
B. Graphs showing morphometric measurements and microCT analysis of bones from *Cog4^+/+^* and *Cog4^+/G512R^* mice (WT, n = 5; Het, n = 4). Horizontal and vertical lines represent the mean with the 95% CI. Statistical analysis was performed using unpaired Student’s t test; ns > 0.05, **= p < 0.01.
C. Graphs showing measurements of long bones and their mineralization ratios for the indicated skeletal elements (WT, n = 4; Het, n = 14; Hom, n = 10). Statistical analysis was performed using Ordinary one-way ANOVA; ns > 0.05, *= p < 0.05, **= p < 0.02, ***= p < 0.001, ****= p < 0.0001.

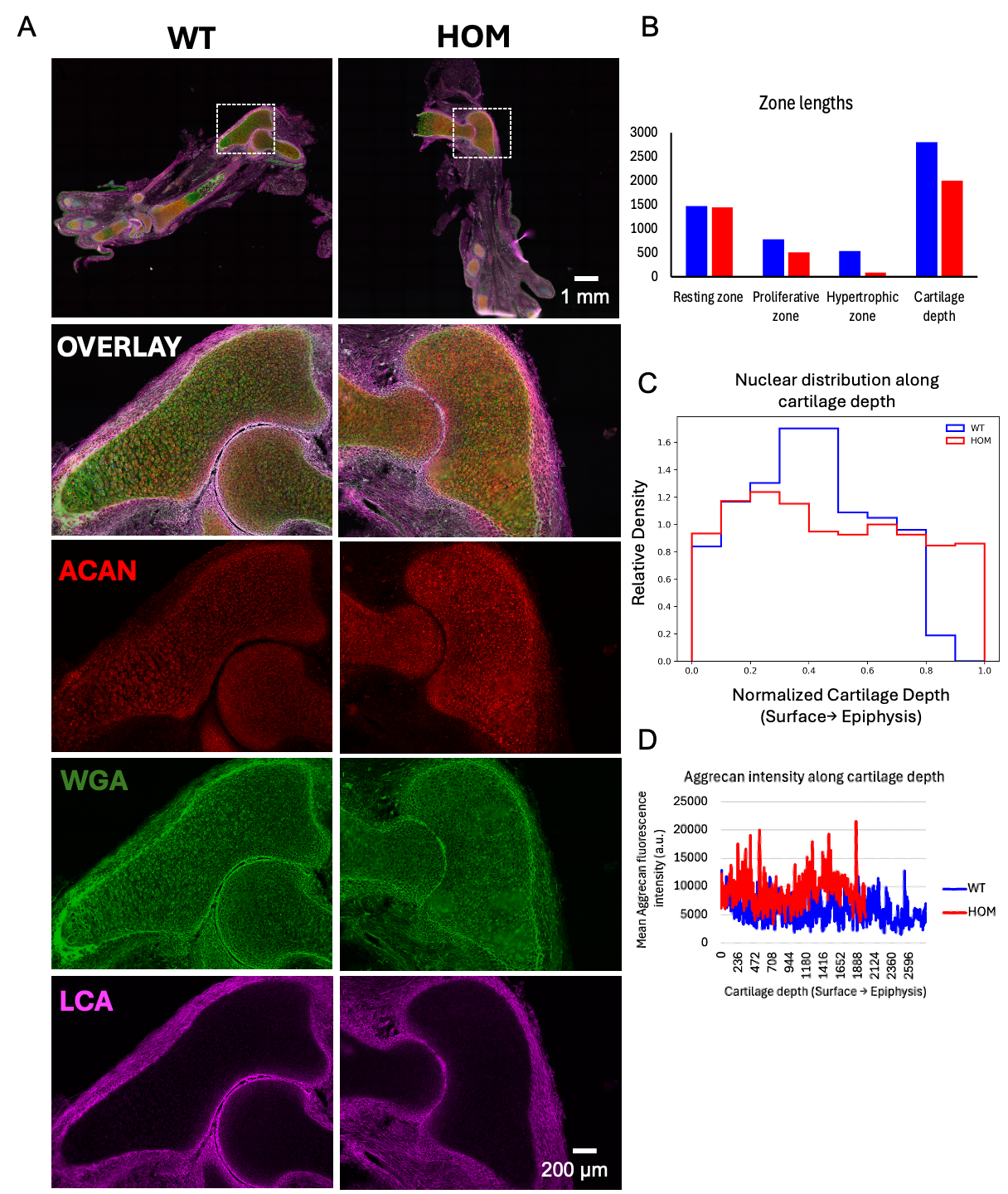

**Figure S2**

A. Representative images of E15.5 mouse embryonic sections highlighting the following markers: Aggrecan, WGA, and LCA. Images are representative of three independent experiments with similar results.
B. Graph showing the lengths of distinct zones in the cartilage shown in zoomed Inset of panel A.
C. Graph representing nuclear distribution along the length of cartilage (shown in the zoomed Inset of panel A) from the surface towards the epiphysis.
D. Graph showing Aggrecan intensity along the length of cartilage shown in the zoomed inset of panel A.

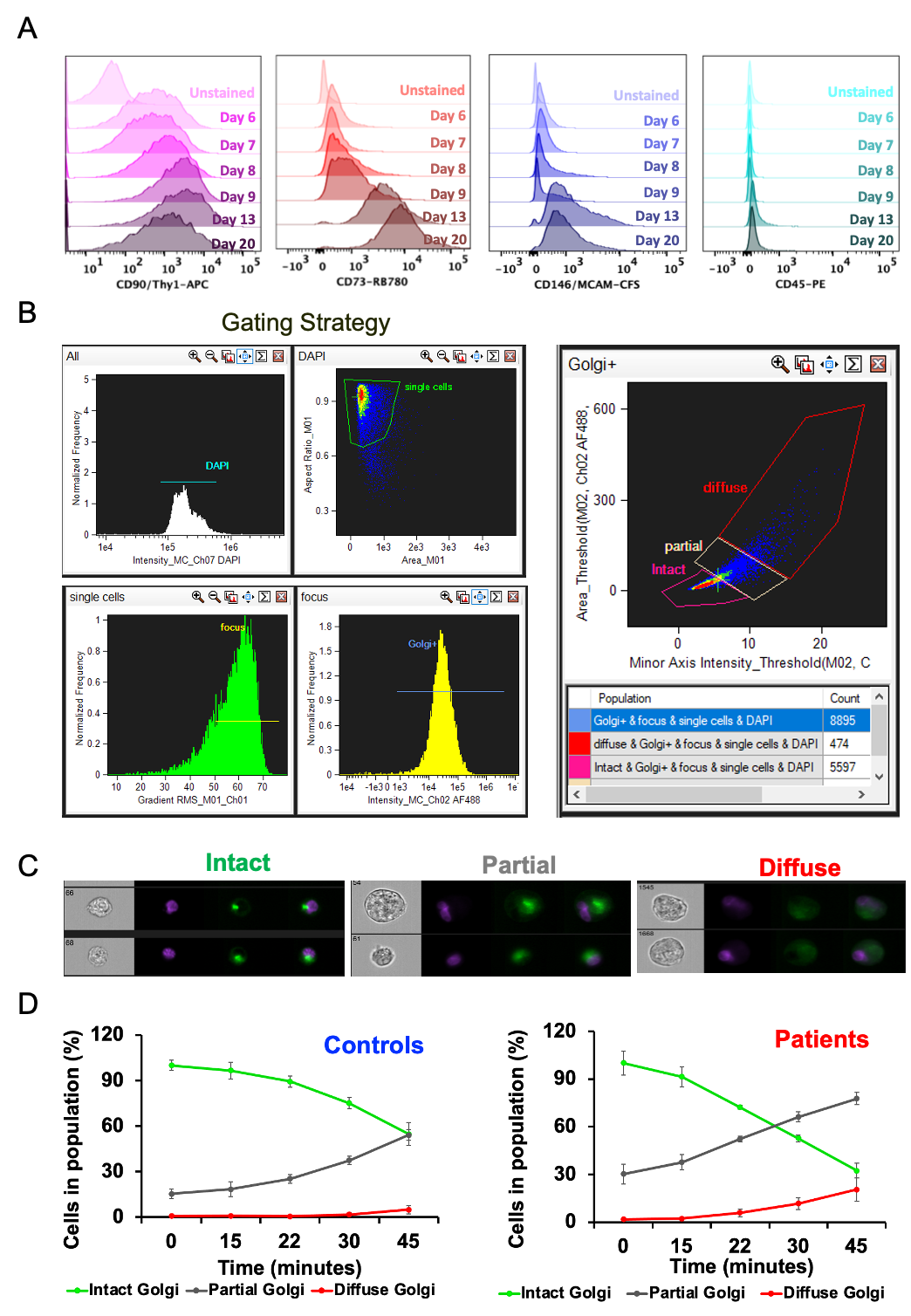

**Figure S3**

A. Characterization of mesenchymal progenitor cells by flow cytometry using mesenchymal markers (CD90, CD73, and CD146) and the negative marker CD45 across different days of differentiation.

B. Gating strategy used to quantify different stages of Golgi diffusion (Intact, Partial, and Diffuse) following BFA treatment.

C. Representative image insets showing the characteristic Golgi morphologies classified as Intact, Partial, and Diffuse.

D. Graph showing quantification of cell populations exhibiting different Golgi morphologies over a 45-minute time course following BFA treatment to assess retrograde trafficking. Error bars represent the standard deviation of three biological replicates.

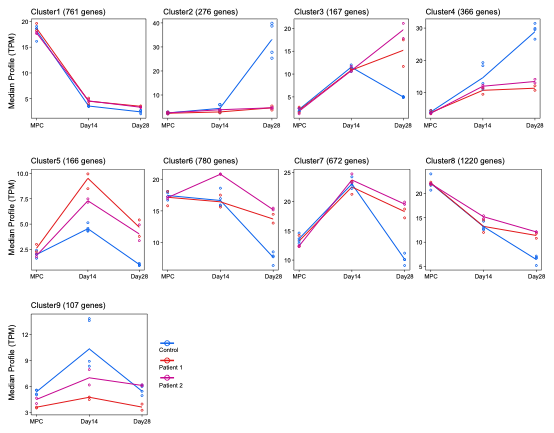

**Figure S4.** Gene expression profile clusters generated using the maSigPro R package to identify groups of genes with similar expression patterns during chondrogenic differentiation.

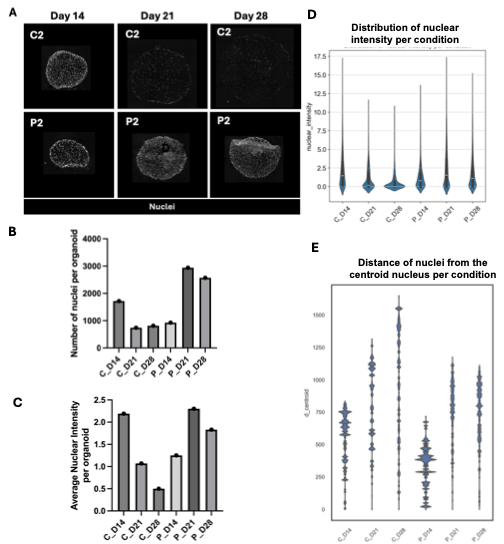

**Figure S5**

A. Representative images of organoid sections showing DRAQ5 signal across different stages of differentiation in controls and patient organoids.

B. Graph showing the number of nuclei across the indicated conditions.

C. Graph showing average nuclear intensity per organoid across the indicated conditions.

D. Graph showing the distribution of nuclear intensity across conditions.

E. Graph showing the distribution of nuclei from the centroid toward the periphery across the indicated conditions.

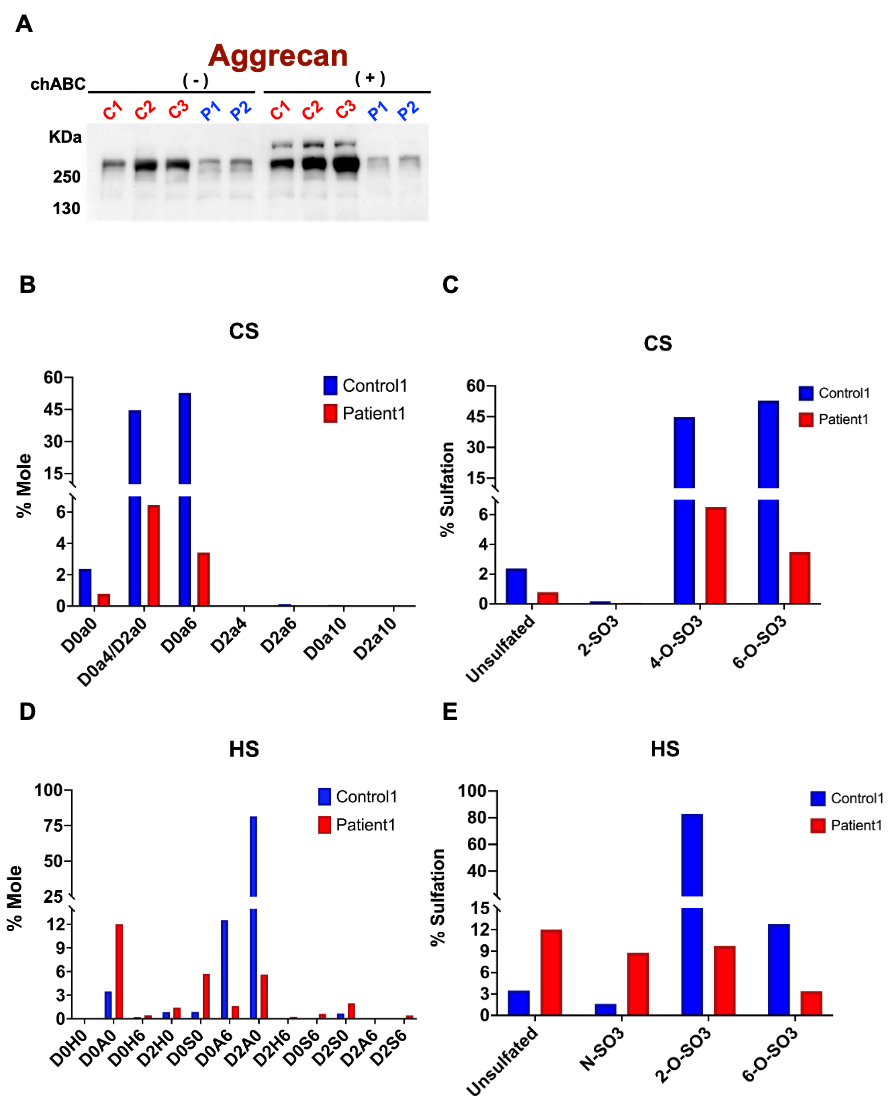

**Figure S6**

A. Representative western blot showing the Aggrecan band in conditioned medium from Day 28 control and patient organoids, before and after chondroitinase treatment.

B. Glycan reductive isotope labeling (GRIL)-LC/MS analysis showing the disaccharide composition of extracted chondroitin sulfate (CS) glycosaminoglycans (GAGs) from Day 28 organoids.

C. Graph showing percentage sulfation in CS GAGs.

D. GRIL-LC/MS analysis showing the disaccharide composition of extracted heparan sulfate (HS) GAGs from Day 28 organoids.

E. Graph showing percentage sulfation in HS GAGs.
